# Resting-state fMRI coherence is selectively diminished around 0.1 Hz, particularly inside watershed areas, in patients with unilateral carotid artery stenosis

**DOI:** 10.1101/2025.06.12.659378

**Authors:** Sangcheon Choi, Gabriel Hoffmann, Sebastian Schneider, Stephan Kaczmarz, Xin Yu, Christine Preibisch, Christian Sorg

## Abstract

In the brain, vasomotor dynamics at infra-slow frequencies (∼0.1 Hz), driven by synchronized oscillations of smooth muscle cells in vessel walls, are thought to play a crucial role in regulating cerebral perfusion and underlie resting-state functional connectivity (FC), typically measured by correlated time courses of functional signals. In particular, rodent studies have demonstrated that vasomotor activity contributes to the coherence of blood oxygenation level dependent (BOLD) signal fluctuations. However, in humans, detecting this contribution non-invasively remains challenging due to the limited spatiotemporal sensitivity of functional magnetic resonance imaging (fMRI) to vasomotion. Given that prior studies have identified internal carotid artery stenosis (ICAS) as an informative conditional lesion model of vasomotor and hemodynamic impairments in humans, we investigated whether ICAS affects interhemispheric BOLD coherence at ∼0.1 Hz. Using a multi-modal fMRI framework integrating resting-state fMRI with quantitative mapping of cerebral blood volume, blood flow, oxygen metabolism, and BOLD time lag, we compared BOLD coherence between patients with asymptomatic unilateral ICAS and healthy controls. Frequency-specific analysis revealed significantly diminished interhemispheric BOLD coherence at ∼0.1 Hz across canonical resting-state networks in ICAS patients, while ultra-slow (<0.05 Hz) coherence remained largely preserved. This reduction was spatially widespread and particularly pronounced in watershed areas, i.e., border zones between major vascular territories, associated with significantly increased lateralization of cerebral blood volume (p < 0.01). Notably, coherence-based FC patterns at ∼0.1 Hz were heterogeneous within watershed areas but homogeneous outside, suggesting an interplay between compensatory mechanisms and cerebrovascular impairment. Taken together, our findings demonstrate that ICAS induces subtle, frequency-and region-specific alterations in interhemispheric FC, consistent with a model in which impaired vasomotor activity impacts on resting-state FC in the human brain.

## INTRODUCTION

Vasomotor activity, arising from synchronized oscillations of smooth muscle cells in vascular walls, has been implicated in changes in arteriole diameter (Nicoll and Webb 1955; Intaglietta 1990; Aalkjær, Boedtkjer, and Matchkov 2011; Wang et al. 2011; Mateo et al. 2017; Drew et al. 2020). These oscillatory vasomotor signals reflect underlying vascular conditions and have been widely observed across species, including humans (Nicoll and Webb 1955; Rosenblum, Bonner, and Oldfield 1987; Seifert, Jager, and Bollinger 1988; Intaglietta 1990; Hudetz, Roman, and Harder 1992; Mayhew et al. 1996; Mitra et al. 1997; Kleinfeld et al. 1998; Obrig et al. 2000; Razavi et al. 2008; Rayshubskiy et al. 2014; Broggini et al. 2024). In animals, invasive techniques such as laser Doppler flowmeter and optical recording have demonstrated that cerebral vasomotor activity typically exhibits slow oscillations peaking around 0.1 Hz within the infra-slow fluctuations (Hudetz, Roman, and Harder 1992; Mayhew et al. 1996; Kleinfeld et al. 1998; Mateo et al. 2017; Broggini et al. 2024; Wang et al. 2011). Importantly, beyond its established role in regulating local cerebral perfusion, vasomotor activity has also been proposed as a physiological basis for functional connectivity (FC) during resting-state (Mateo et al. 2017). This hypothesis is primarily supported by evidence that spontaneous vasomotor oscillations, synchronized across homotopic cortical regions (i.e., symmetric regions in both hemispheres) and induced by ongoing neuronal activity, contribute to coherent blood oxygenation dependent level (BOLD) functional magnetic resonance imaging (fMRI) signals that underlie resting-state FC in awake rodents as demonstrated by invasive optical imaging (Mateo et al. 2017; Drew et al. 2020; Broggini et al. 2024). Coherence-based analysis has further demonstrated that spontaneous arteriole diameter oscillations, driven by vasomotion (∼0.1 Hz) in the infra-slow fluctuations, can modulate BOLD fMRI signal coherence, particularly in homotopic cortical regions (Mateo et al. 2017; Drew et al. 2020). Notably, rodent fMRI studies have reported a distinct peak in power spectral density (PSD) near 0.1 Hz in somatosensory cortices, suggesting that a contribution of vasomotion-related mechanisms to inter-hemispheric FC (Williams et al. 2010; Choi et al. 2023).

However, in humans, only a few invasive studies have reported spatially distinct vasomotor oscillations across the cortex at the vasomotor frequency (∼0.1 Hz) under intraoperative conditions (Rosenblum, Bonner, and Oldfield 1987; Rayshubskiy et al. 2014). Despite these observations, evidence linking vasomotor activity to resting-state BOLD FC in humans using non-invasive fMRI remains scarce due to high signal variability and overlap with Mayer waves in the same frequency band (Julien 2006; Elstad et al. 2011; Mitra et al. 1997; Razavi et al. 2008; Rayshubskiy et al. 2014). Vasomotor activity, which regulates the arterial reservoir volume (Nicoll and Webb 1955), is considered ubiquitous in vascular structures (Broggini et al. 2024) and may serve as a potential index of cerebrovascular disease. Indeed, disrupted vasomotor activity has been observed in both high-risk stroke mouse models (Zhang et al. 2024) and in patients with internal carotid artery stenosis (ICAS) (Silvestrini et al. 1996; Gur, Bova, and Bornstein 1996; Silvestrini et al. 2000; Gur and Bornstein 2006; Avirame et al. 2015). Given the evidence of vasomotor impairment in patients with ICAS, it is plausible that inter-hemispheric resting-state (rs-) fMRI coherence, particularly in the ∼0.1 Hz frequency band, may be altered in patients with ICAS and could serve as a frequency-specific signature of cerebrovascular disease based on impaired vasomotor activity. To date, no direct model of vasomotor dynamics for FC currently exists in humans. Although prior studies have suggested that ICAS can be considered a natural lesion model of vasomotion (Silvestrini et al. 1996; Gur, Bova, and Bornstein 1996; Silvestrini et al. 2000; Gur and Bornstein 2006; Avirame et al. 2015), its impact is not limited to vasomotion alone. ICAS is also associated with hemodynamic impairments (i.e., perfusion alterations) (Gleissner et al. 2022; Kaczmarz et al. 2021; Kaczmarz et al. 2018; Gottler et al. 2019; Schneider et al. 2022; Schneider et al. 2023) that influence BOLD FC. Therefore, ICAS represents an informative conditional lesion model for investigating alterations in cerebrovascular regulation that may involve impaired vasomotion in humans.

In addition, multi-parametric hemodynamic MRI studies of patients with asymptomatic unilateral ICAS have revealed unilateral vascular-hemodynamic impairments in cerebral blood flow (CBF) and cerebral blood volume (CBV) (Gleissner et al. 2022; Kaczmarz et al. 2021; Kaczmarz et al. 2018; Gottler et al. 2019; Schneider et al. 2022; Schneider et al. 2023), while the cerebral metabolic rate of oxygen (CMRO_2_) remains largely preserved, suggesting intact neuronal function (Gottler et al. 2020). The most pronounced alterations in CBF and CBV have been observed in watershed areas, i.e., cortical and internal border zones between major perfusion territories that are particularly vulnerable to hypoperfusion (Kaczmarz et al. 2021), suggesting that vasomotor activity may be reflected in macroscopic CBV measures (He et al. 2018; Holstein-Ronsbo et al. 2023; Kilic and Devor 2023). In addition, disrupted coupling between CBF and CMRO_2_ has been reported in the ICAS patients, indicating altered flow-metabolism dynamics (Gottler et al. 2019). Although correlation-based rs-fMRI analyses have shown that vasomotor impairment is strongly associated with decreased FC in patients with high-grade ICAS (Avirame et al. 2015), minimal or no group-level differences in homotopic correlation between healthy controls and ICAS patients (after controlling for hemodynamic impairments) suggest that vascular-hemodynamic impairments may influence resting-state FC in more subtle or complex ways (Schneider et al. 2022; Schneider et al. 2023).

Taken together, previous studies suggest that inter-hemispheric coherence of rs-fMRI signals in patients with asymptomatic unilateral ICAS may be altered in both frequency-specific and spatially dependent ways. We therefore expect that in ICAS patients, hemodynamic and vasomotor-related impairments of cerebrovascular regulation are associated with spectrally and spatially specific alterations in inter-hemispheric resting-state BOLD FC. Furthermore, we propose that frequency-resolved coherence analysis can uncover subtle but meaningful disruptions in brain networks that may not be detectable using conventional correlation methods, thereby providing new insights into the physiological basis of FC in the human brain. Our investigation is guided by three central specific hypotheses regarding the impact of unilateral ICAS: 1) Frequency specificity: We hypothesize that the most prominent alterations in BOLD signal oscillations and coherence will occur in the ∼0.1 Hz frequency range (particularly 0.08-0.1 Hz (Mitra et al. 1997; Rayshubskiy et al. 2014; Obrig et al. 2000)), which has been associated with vasomotion-related processes (Avirame et al. 2015; Mateo et al. 2017), whereas ultra-slow coherence below 0.05 Hz, reflecting neuronal activity (He et al. 2018) and arousal brain state (Raut et al. 2021) will remain largely preserved. 2) Brain-wide effect: We hypothesize that disruptions in ∼0.1 Hz-specific inter-hemispheric coherence will be spatially widespread across the whole brain, supporting further the vascular contribution, rather than neuronal origin, to FC alterations (Mitra et al. 1997; Razavi et al. 2008; Broggini et al. 2024; Lambers et al. 2023). 3) Regional vulnerability: We hypothesize that disrupted BOLD signal fluctuations, inter-hemispheric coherence alterations, and corresponding changes in vasomotion-related hemodynamic parameters, particularly CBV, will be concentrated in vulnerable vascular territories such as watershed areas, which have been previously demonstrated to be most affected by hypoperfusion (Torvik 1984; Bilecen et al. 2002; Kaczmarz et al. 2021).

In this study, we combined rs-fMRI with quantitative hemodynamic MRI to investigate how asymptomatic unilateral ICAS alters frequency-specific inter-hemispheric BOLD signal coherence and its coupling to cerebral perfusion parameters. To test our hypotheses, we developed a novel analytic framework integrating rs-fMRI-based spectral PSD and coherence analysis with multi-parametric MRI maps including relative CBV, CBF, relative CMRO₂, and BOLD time lag (defined as relative time lag of cortical regions to the sagittal superior sinus). This framework was applied to both healthy controls and patients with unilateral ICAS, particularly inside watershed areas as well as across large-scale resting-state brain networks. To assess frequency specificity (hypothesis 1), we compared inter-hemispheric PSD and coherence between healthy controls and patients with asymptomatic unilateral ICAS across distinct frequency bands (<0.05 Hz and ∼0.1 Hz) and across both homotopic and heterotopic regions of interest (ROI) pairs. To assess brain-wide effects (hypothesis 2), we examined inter-hemispheric coherence patterns across intrinsic resting state networks defined by Yeo and colleagues (Yeo et al. 2011). To assess regional vulnerability (hypothesis 3), we compared inter-hemispheric PSD, coherence, and quantitative hemodynamic parameters between distinct perfusion territories (Seifert, Jager, and Bollinger 1988), specifically regions inside versus outside watershed areas, i.e., vascular border zones that are particularly vulnerable to reductions in CBF in patients with ICAS (Kaczmarz et al. 2021).

## MATERIALS and METHODS

In this study, we analyzed a subsample of existing multi-modal MRI data from our previous study, from which different aspects have already been published (Gottler et al. 2020; Gleissner et al. 2022; Kaczmarz et al. 2021; Kaczmarz et al. 2018; Gottler et al. 2019; Schneider et al. 2022; Schneider et al. 2023).

## SUBJECTS AND MULTI-MODAL MRI DATA ACQUISITION

### Subjects

All participants provided written informed consent prior to scanning in accordance with the policies of the institution’s Human Research Committee. The study was approved by the Institutional Review Board of the Klinikum rechts der Isar at Technical University of Munich. Patients with unilateral asymptomatic ICAS were recruited through the outpatient clinic for carotid stenoses at the Department of Vascular and Endovascular Surgery, while healthy control participants were recruited via word-of-mouth advertisement. As previously reported (Gottler et al. 2020; Gleissner et al. 2022; Kaczmarz et al. 2021; Kaczmarz et al. 2018; Gottler et al. 2019; Schneider et al. 2022; Schneider et al. 2023), the study enrolled 29 patients with high-grade, unilateral, asymptomatic ICAS (confirmed by duplex ultrasonography; all >70% stenosis based on NASCET criteria (NASCET 1991)) and 30 age-matched healthy controls. Each participant underwent a clinical assessment that included a medical history review, basic neurological and psychiatric screening, and an MRI scan. Patients were classified as clinically asymptomatic if they exhibited no neurological symptoms such as stroke, transient ischemic attack (TIA), or paresis. Individuals with a history of neurological or psychiatric disorders, clinically significant structural brain abnormalities, or MRI contraindications were excluded. Due to insufficient MRI data quality, two patients and four healthy controls were excluded from the analysis, resulting in a final sample of 27 patients (11 females; mean age 70.3 ± 7.2 years) and 26 healthy controls (15 females; mean age 70.3 ± 4.8 years). Detailed demographic information was described in Supplementary Information (**Table S4**).

### Multi-modal rs-fMRI and multiparametric quantitative MRI data acquisition

All MRI data were acquired on a whole-body 3T Ingenia MRI scanner (Philips Healthcare, Best, The Netherlands) equipped with a 16-channel (dynamic susceptibility contrast (DSC) MRI) or a 32-channel head coil (for others) (**Fig. 1A**). Detailed acquisition parameters are described in the Supplementary Information. In short: For assessing PSD, coherence of BOLD fMRI signal time series, T2*-weighted multiband echo planar imaging (EPI) data were acquired under resting-state conditions. The acquisition parameters were as follows: repetition time (TR) = 1200 ms, echo time (TE) = 30 ms, total acquisition time = 10 minutes, flip angle (FA) = 70°, multiband factor = 2, SENSE factor = 2, 38 slices, matrix size = 192x192, voxel size = 3x3x3 mm^3^, and 500 dynamic scans. For the anatomical reference (**Fig. 1B**), T1-weighted magnetization prepared rapid gradient echo (MPRAGE) data were acquired with 170 sagittal slices (**Fig. 1C**).

**Figure 1.**
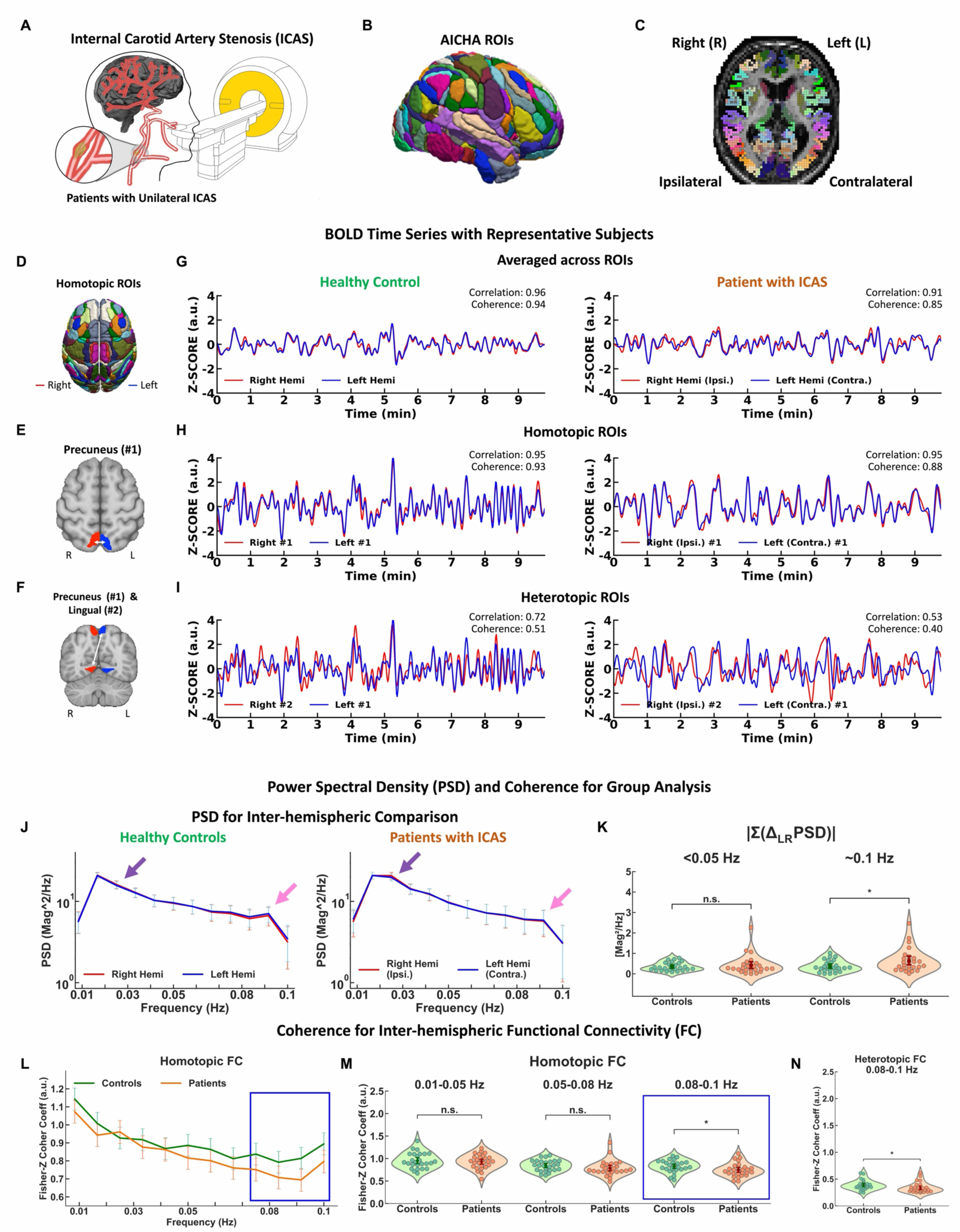
Frequency dependence of homotopic and heterotopic FC in healthy controls and patients with ICAS. **A.** Conceptual illustration of multi-modal MRI measurements in asymptomatic patients with unilateral ICAS at 3T. **B.** Surface representation of AICHA brain parcellation (N = 384 ROIs). **C.** AICHA ROIs displayed on an axial T1-weighted MPRAGE image; identical colors indicate homotopic regions. **D-I**. BOLD time series for one representative subject from each group. **D.** illustration of homotopic ROI pairs defined by AICHA atlas. **E** and **F.** AICHA ROIs corresponding to a representative homotopic ROI pair (**E**; Precuneus) and a heterotopic ROI pair (**F**; Precuneus and Lingual). Red and blue indicate right and left hemispheric regions, respectively. **G.** Averaged Z-score normalized BOLD signal time series from right (red) and left (blue) hemispheres in a representative healthy control (left) and patient with ICAS (right). Correlation and coherence values are shown in each panel (top right). **H** and **I.** Z-score normalized BOLD time series from the homotopic (**H**) and heterotopic (**I**) ROI pairs for the same representative subjects as in panel **G**. Correlation and coherence values are shown in each panel. **J-N**. Power spectral density (PSD) and coherence for group analysis. **J.** Group-averaged PSD curves of intra-hemispheric BOLD signals from healthy controls (left, n=26) and patients with ICAS (right, n=27). Error bars indicate mean ± SD across subjects. Purple and pink mark spectral peaks at ∼0.02 Hz and ∼0.1 Hz, respectively. **K.** Group-level absolute summed PSD differences between the left and right hemispheres, |Σ(Δ_LR_ PSD)| at <0.05 Hz and ∼0.1 Hz (0.08-0.1 Hz, see **Methods**) for healthy controls (green) and patients with ICAS (orange). **L** and **M.** Coherence for inter-hemispheric FC (healthy controls: green, patients with ICAS: orange). **L.** Group-level inter-hemispheric coherence (homotopic FC) at the infra-slow frequency band (0.01-0.1 Hz). The blue box marks the frequency range of ∼0.08-0.1 Hz. Error bars indicate mean ± SEM across subjects. **M.** Comparison of homotopic FC across three distinct frequency bands: 0.01-0.05 Hz, 0.05-0.08 Hz, and 0.08- 0.1 Hz (highlighted by blue box). **N.** Heterotopic FC at ∼0.08-0.1 Hz. Statistical significance based on two-sample t-tests (*: p-value < 0.05; n.s.: not significant). Diamond markers and outlines represent group means and 95% confidence intervals, respectively.

Quantitative perfusion and oxygenation parameter were acquired and processed as described in detail previously (Gottler et al. 2019; Kaczmarz et al. 2021; Schneider et al. 2023). Relative CBV was derived from DSC MRI data following bolus injection of gadolinium-based contrast agent (Gd-Dota; concentration 0.5 mmol/ml, dose 0.1 mmol/kg, minimum 7.5 mmol per subject). CBF was measured using pseudo-continuous arterial spin labeling (pCASL) sequence in accordance with the ISMRM Perfusion Study Group consensus recommendations (Alsop et al. 2015) using a prolonged post-labeling delay of 2000 ms and applying background suppression. The oxygen extraction fraction (OEF) was calculated from R2′ (= 1⁄*T*2^∗^ − 1/*T*2) and CBV, where *T*2^∗^and *T*2 were derived from separate GRE and GRASE sequences, respectively, according to the multiparametric quantitative BOLD approach (Kaczmarz, Hyder, and Preibisch 2020). Relative CMRO_2_ was then estimated from the product of relative OEF and CBF (Rodgers, Detre, and Wehrli 2016; Biondetti, Cho, and Lee 2023).

## MRI DATA PROCESSING

All data processing was performed using MATLAB (Mathworks, Natick, MA, version R2023a), Python (version 3.9.13), and R (version 4.4.0). Detailed descriptions of the processing and analysis workflows have been published in previous studies (Choi et al. 2023; Choi et al. 2022; Schneider et al. 2022; Schneider et al. 2023). All figures were generated using Python (version 3.9.13), R (version 4.4.0), and FreeSurfer (version 7.4.1).

## DATA PROCESSING FOR BOLD-fMRI

### BOLD signal processing

We analyzed a subsample of 53 participants—27 patients (11 female) and 26 healthy controls (15 females)—selected based on sufficient MRI for analysis (e.g., absence of motion artifacts and valid values in Atlas of Intrinsic Connectivity of Homotopic Areas (AICHA) ROIs).

As described in previous works (Schneider et al. 2022; Schneider et al. 2023), BOLD fMRI data were pre-processed using the Data Processing Assistant for Resting-State fMRI advanced edition (DPARSFA) toolbox V5.0_200401 (Chao-Gan and Yu-Feng 2010) and SPM12 (v7771; Wellcome Trust Centre for Neuroimaging, UCL, London, UK) using MATLAB (R2019b; MathWorks, Natick, MA, USA). Pre-processing included discarding of the first ten EPI volumes to reach steady-state magnetization, spatial realignment of acquired EPI volumes, slice-timing correction, reorientation to the anterior-posterior commissure, nuisance covariate regression (six motion parameter and a linear trend), spatial smoothing with a 6 mm full width at half maximum kernel, and temporal bandpass filtering (0.01 to 0.1 Hz). Anatomical images were first neck cropped and then co-registered with functional EPI images, followed by segmentation using Diffeomorphic Anatomical Registration Through Exponentiated Lie Algebra (DARTEL) (Ashburner 2007).

For each individual subject, EPI fMRI data were parcellated using the AICHA (Joliot et al. 2015) (**Fig. 1B**), which resulted in 192 pairs of homotopic ROIs, yielding a total of 384 cortical and subcortical gray matter ROIs (**Fig. 1C**).

BOLD time series were extracted from the 192 homotopic ROI pairs, and Z-score normalization was applied prior to PSD, coherence, and correlation analyses. Z-score normalization was computed as *z* = (*x* − μ)/σ, where *x* represents the original fMRI time series and μ and σ denote its mean and the standard deviation, respectively (using the ‘zscore’ function in MATLAB). In representative cases—a healthy control and a patient with unilateral ICAS (**Fig. 1G-I**)—Z-score normalized fMRI time series were averaged across homotopic ROI pairs to illustrate differences in temporal patterns (**Fig. 1D and G**). To more specifically compare infra-slow oscillations (<0.1 Hz), representative Z-score normalized fMRI time series were also extracted from one homotopic ROI pair (precuneus) and one heterotopic ROI pair (precuneus and lingual gyrus) in both subjects (**Fig. 1H and I**).

### PSD analysis

Z-score normalized fMRI time series were transformed into PSD values to assess amplitude differences in resting-state BOLD signal power between the two groups. For group analysis, PSDs of bilateral rs-fMRI signals were compared in the typical resting-state frequency band, i.e., 0.01-0.1 Hz (Fox and Raichle 2007; Cordes et al. 2001; He et al. 2018; Choi et al. 2022; Choi et al. 2023; Schneider et al. 2022; Schneider et al. 2023) (**Fig. 1J**). To quantify inter-hemispheric differences, PSD differences between the left and right hemispheres were summed across all homotopic ROI pairs and converted to absolute values, |Σ(Δ_LR_ PSD)| =|Σ(PSD_Left_ - PSD_Right_|, at two distinct frequency bands; 0.01-0.05 Hz and 0.08-0.1 Hz (**Fig. 1K**). PSD estimation was performed using Welch’s method (‘pwelch’ function in MATLAB) with a Hamming window length of 128, Fast Fourier transform (FFT) length of 100, and 50% overlap. The frequency resolution was 1/120 s = 0.0083 Hz. The two distinct frequency bands were 0.01-0.05 Hz (referred to as <0.05 Hz) and 0.08-0.1 Hz (referred to as ∼0.1 Hz in the following).

### Spectral coherence analysis

Coherence analysis was employed to investigate the spectral dynamics of inter-hemispheric FC, as coherence is widely regarded as a frequency-domain measure of FC (He et al. 2018; Choi et al. 2022; Choi et al. 2023; Sun, Miller, and D’Esposito 2004; Yaesoubi et al. 2015). Higher coherence coefficients are generally interpreted as indicating stronger FC, whereas lower coefficients suggest reduced FC. In this study, spectral coherence was treated as a key marker for frequency-specific inter-hemispheric FC. To examine differences in coherence patterns between healthy controls and patients with unilateral ICAS, group-averaged spectral coherence was analyzed for both homotopic and heterotopic FC across the infra-slow oscillatory frequency range (<0.1 Hz), specifically focusing on two distinct frequency bands: <0.05 Hz and ∼0.1 Hz. Z-score normalized fMRI time series were used for coherence calculations to eliminate potential bias stemming from the baseline signal intensity differences between two hemispheres.

The coherence coefficient between two signals *x* and *y*—representing either homotopic or heterotopic ROI pairs—is defined as follows:

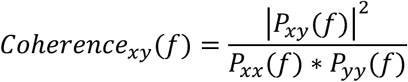

where *P_xx_*(*f*) and *P_yy_*(*f*) denote the auto spectral densities of signals *x* and *y*, respectively, and *P_xy_*(*f*) is the cross-spectral density between them.

Coherence was computed using the magnitude-squared coherence estimate method (‘mscohere’ function in MATLAB, Mathworks, Natick, MA) with a hamming window length: 128, FFT length: 100, overlap: 50%). As in the PSD analysis, the frequency resolution was 1/120 s = 0.0083 Hz, allowing for the characterization of coherence patterns across the infra-slow oscillatory frequency range (<0.1 Hz). Two distinct frequency bands were examined: 0.01-0.05 Hz (referred to as <0.05 Hz) and 0.08-0.1 Hz (referred as to ∼0.1 Hz). To facilitate group-level comparison, coherence coefficients between inter-hemispheric ROIs were Fisher Z-transformed prior to averaging within each frequency bands and group (Corey, Dunlap, and Burke 1998) (**Fig. 1L-N**).

### Temporal correlation analysis

To complement the coherence analysis, temporal correlation analysis was conducted to assess inter-hemispheric FC across homotopic and heterotopic ROI pairs. As in the coherence analysis, Z-score normalized fMRI time series were used for both healthy controls and patients with unilateral ICAS.

The Pearson correlation coefficient between two signals *x* and *y* is defined as the following equation:

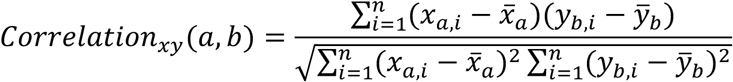

where *x* and *y* are the fMRI time series from ROIs *a* and *b*, respectively; *i* indicates the time points; and 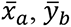 are the mean values of each time series.

Correlation coefficients were computed using ‘corr’ function in MATLAB (type: Pearson, rows: pairwise). As with coherence, Fisher-Z transform was applied to correlation values prior to averaging across inter-hemispheric ROI pairs within each group (**Fig. S5 A**).

### Network-specific FC analysis

To assess the regional specificity of inter-hemispheric rs-fMRI, seven canonical rs-fMRI networks were utilized as defined by Yeo and colleagues (Yeo et al. 2011). These networks were derived from a dataset of 1,000 healthy individuals scanned on 3T Tim Trio MRI scanners (Siemens, Erlangen, Germany), and are widely used to characterize large-scale distributed network organization in human fMRI research. The seven networks included: Default Mode Network (DMN), Somatomotor Network (SMN), Dorsal Attention Network (DAN), Visual Network (VIN), Salience Network (SAN), Limbic Network (LIN), and Executive Control Network (ECN). In addition, subcortical network (SubCort) was delineated.

For network-specific FC analysis, inter-hemispheric coherence matrices were computed using homotopic and heterotopic AICHA ROI pairs across the infra-slow oscillatory frequency band (0.01-0.1 Hz). Each AICHA ROI was assigned to one of the canonical rs-fMRI networks based on spatial overlap with the AICHA atlas (Joliot et al. 2015) and Yeo’s network parcellation (Yeo et al. 2011), following procedures described by Labache et al. (Labache et al. 2023) (**Fig. 2A**). The coherence matrices were then reordered for each participant group (healthy controls and patients with unilateral ICAS) according to the canonical networks order: VIN, SMN, DAN, SAN, LIN, ECN, DMN, and SubCort. The reordered coherence matrices were averaged across three infra-slow frequency bands: the full infra-slow band (0.01-0.1 Hz; **Fig. 2B**), a ultra-slow oscillatory frequency band (0.01-0.05 Hz; **Fig. 2C**), and a slow oscillatory frequency band (0.08-0.1 Hz; **Fig. 2E**). Fisher-Z transform was applied prior to averaging and inverted after averaging to ensure appropriate statistical treatments. To assess regional specificity of inter-hemispheric FC across the networks, coherence values were averaged within each network for the two distinct frequency bands (<0.05 Hz and ∼0.1 Hz) (**Fig. 2D and F**). Detailed information on the mapping of AICHA ROIs to canonical networks is provided in the Supplementary **Table S1**).

**Figure 2.**
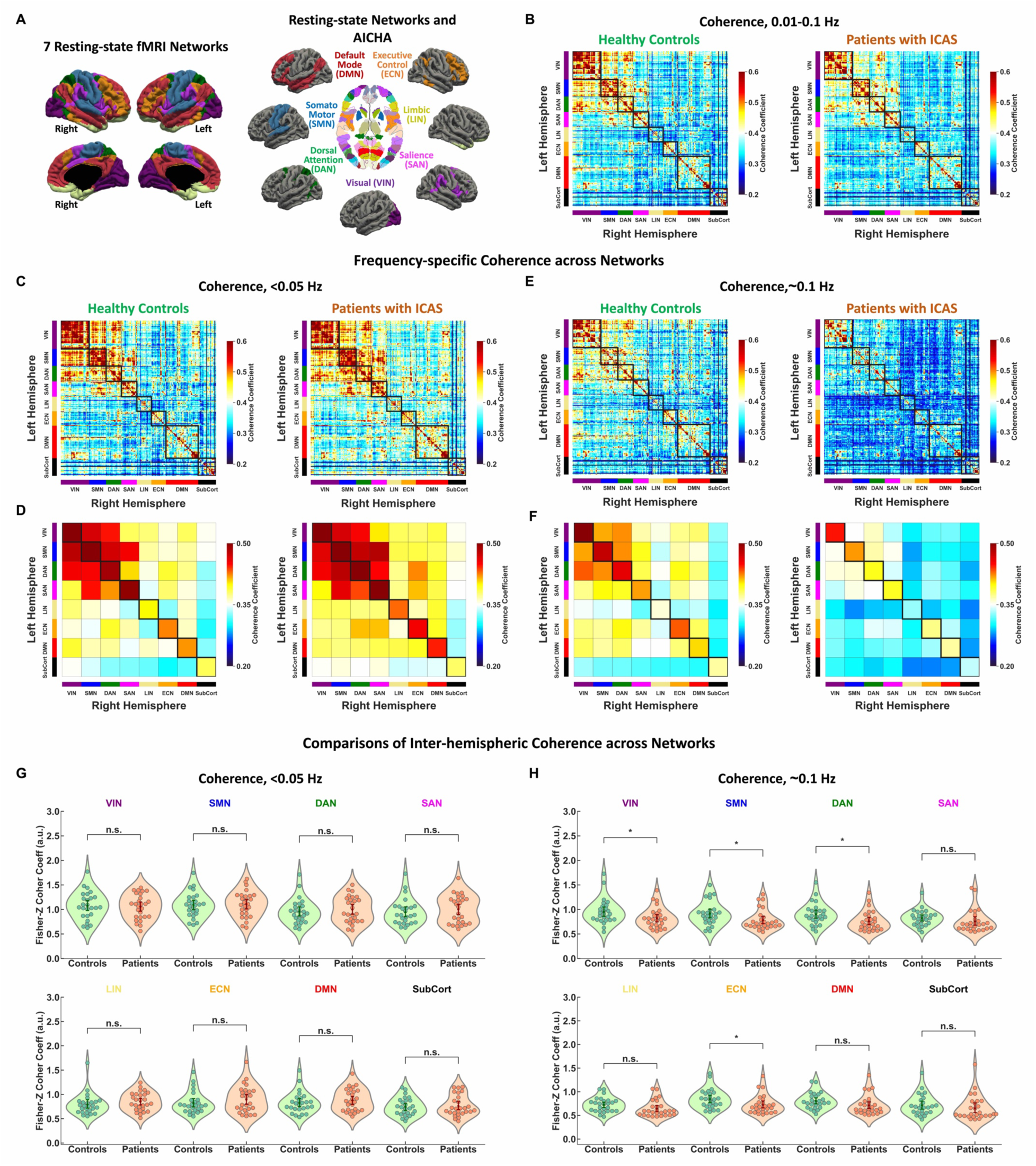
Frequency-dependent FC across networks in healthy controls and patients with ICAS. **A.** Surface representation of seven canonical rs-fMRI networks in right and left hemispheres (left), and their correspondence to AICHA ROIs (right). Overlapping AICHA ROIs were assigned to individual networks: Default Mode Network (DMN, red), Somatomotor Network (SMN, blue), Dorsal Attention Network (DAN, green), Visual Network (VIN, purple), Salience Network (SAN, magenta), Limbic Network (LIN, cream), and Executive Control Network (ECN, orange). The subcortical network (SubCort, black) was also included. **B-F.** Inter-hemispheric coherence matrices of reordered AICHA ROIs across networks for healthy controls (left) and patients with ICAS (right). **B.** Coherence matrices for the full infra-slow frequency range (0.01-0.1 Hz). **C** and **D.** Frequency-specific coherence analysis at <0.05 Hz (0.01-0.05 Hz): **C.** Full network-level coherence matrices. **D.** Network-averaged coherence matrices across the eight networks. **E** and **F.** Frequency-specific coherence analysis at ∼0.1 Hz (0.08-0.1 Hz): **E.** Full network-level coherence matrices. **F.** Network-averaged coherence. **G** and **H.** Group comparisons of inter-hemispheric coherence across networks between healthy controls (green) and patients with ICAS (orange): **G.** At <0.05 Hz. **H.** At ∼0.1 Hz. Statistical comparisons were performed using two-sample *t*-tests (n.s.: no significance and *: p-value<0.05). Diamond markers and outlines represent group means and 95% confidence intervals.

### Identification of distinct perfusion territory

In patients with ICAS, the most vulnerable brain regions are located in the watershed areas, as these regions are particularly susceptible to reductions in CBF. Watershed areas are located at the cortical border zones between major arterial territories—namely, of the anterior, middle, and posterior cerebral arteries (ACA, MCA, and PCA)—as well as at internal border zone between lateral lenticulo-striate arteries (LSA) and MCA (Torvik 1984; Momjian-Mayor and Baron 2005; Kaczmarz et al. 2021) (**Fig. 3A**). Because these regions are furthest from primary arterial supply and depend on distal branches of the major arteries, watershed areas can be defined based on temporal perfusion delay (Kaczmarz et al. 2018).

**Figure 3.**
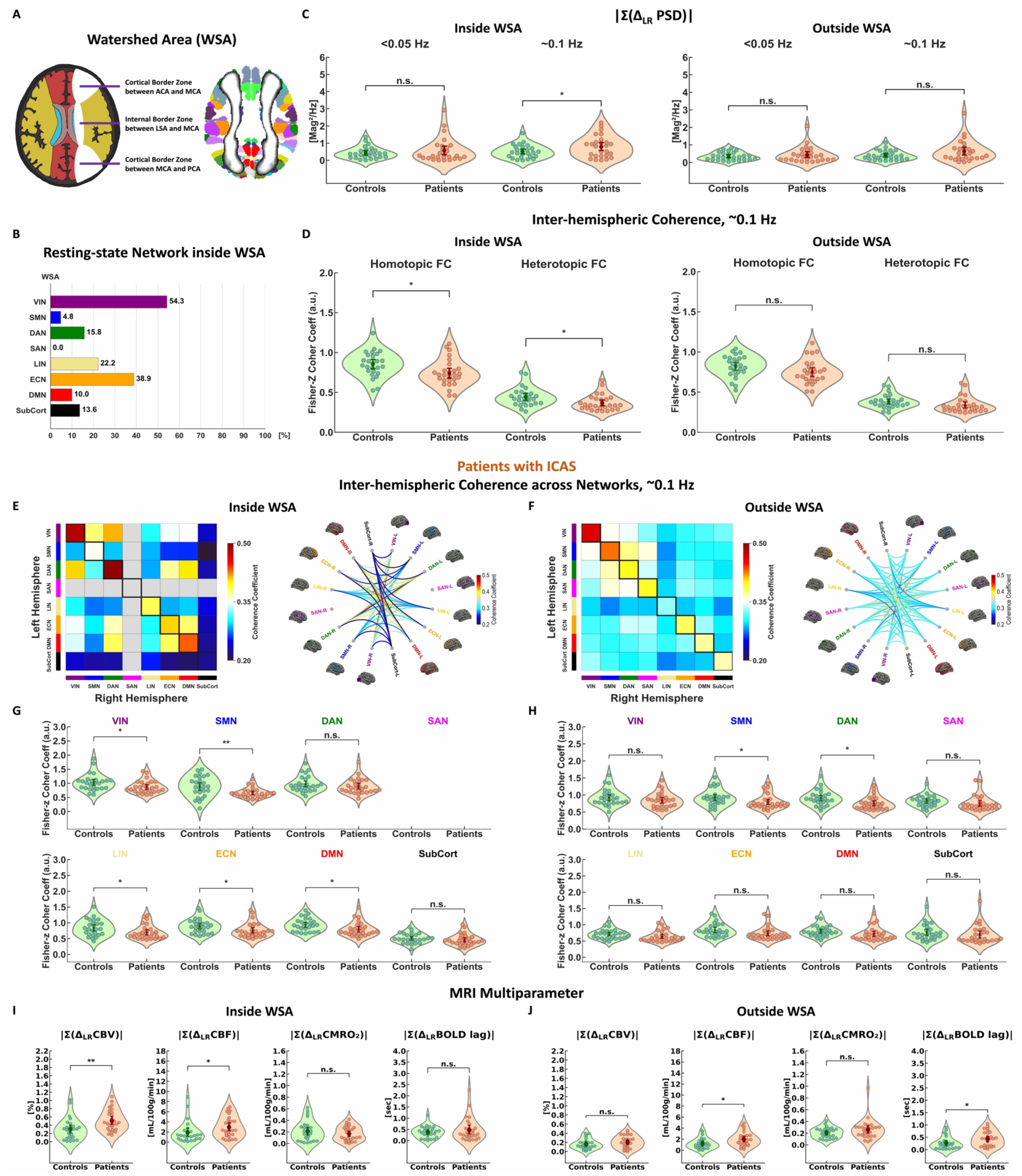
Frequency dependent FC inside and outside of watershed areas. **A.** Schematic illustration of watershed areas (WSAs, white), i.e., border zones: the cortical border zone between the anterior and middle cerebral arteries (ACA–MCA, top), the internal border zone between the lateral striate arteries (LSA) and MCA (middle), and the cortical border zone between the middle and posterior cerebral arteries (MCA–PCA, bottom). Right: Group-average bilateral WSA mask (white) from ICAS patients overlaid on the AICHA atlas (colored ROIs indicate homotopic regions). **B.** Proportional distribution of AICHA ROIs within WSAs across resting-state networks and subcortical regions: VIN (visual network, purple): 54.3%, SMN (somatomotor network, blue): 4.8%, DAN (dorsal attention network, green): 15.8%, SAN (salience network): 0%, LIN (limbic network, cream): 22.2%, ECN (executive control network, orange): 38.9%, DMN (default mode network, red): 10.0%, and SubCort (subcortical regions, black): 13.6%. **C** and **D.** Group-level comparisons of inter-hemispheric PSD and coherence values inside and outside WSAs for healthy controls (green) and patients with ICAS (orange). **C.** Absolute inter-hemispheric PSD differences |Σ(Δ_LR_ PSD)| at <0.05 Hz and ∼0.1 Hz (0.08-0.1 Hz). **D.** Inter-hemispheric coherence values for homotopic and heterotopic FCs inside (left) and outside WSAs (right). **E-H.** Network-specific inter-hemispheric coherence at ∼0.1 Hz (0.08-0.1 Hz) in ICAS patients: **E** and **F.** Network-wise coherence matrix (left) and corresponding circular visualizations (right) inside (**E**) and outside (**F**) WSAs. Gray color in the SAN row and column (**E**) denotes unassigned values. **G** and **H.** Group comparisons of inter-hemispheric coherence across networks inside (**G**) and outside (**H**) WSAs. Two-sample t-tests were used to assess group differences (*: p-value<0.05, n.s.: not significant). **I** and **J.** Group comparisons of inter-hemispheric differences in quantitative MRI parameters (CBV, CBF, CMRO₂, and BOLD time lag) inside (**I**) and outside (**J**) WSAs. Values represent absolute sums of left–right hemisphere differences (|Σ(Δ_LR_ MRI Parameter)|). Wilcoxon rank-sum tests were used to assess statistical significance (*: p-value < 0.05, **: p-value < 0.01, and n.s.: no significance). Diamond markers and outlines indicate group means and 95% confidence intervals.

In this study, individual watershed areas were identified using DSC MRI-derived time-to-peak (TTP) maps, following the method described by Kaczmarz and colleagues (Kaczmarz et al. 2018; Kaczmarz et al. 2021). To alleviate variability in the definition and boundaries of watershed areas and to ensure spatial alignment with AICHA ROIs, a group-level watershed area mask was generated by averaging the individual masks across subjects (n = 52; one patient was excluded due to insufficient data quality) Based on the spatial overlap between the AICHA atlas and the group-defined watershed regions, AICHA ROIs were assigned to the watershed territory using a threshold of 95% overlap. A total of 41 ROI pairs, spanning canonical rs-fMRI networks and subcortical regions, were identified and confirmed though additional visual inspection. The watershed area mask was computed using visualization tools from Nilearn Python library (https://nilearn.github.io/dev/modules/plotting.html) and array transformation functions from Numpy Python library (https://numpy.org).

### Spectral PSD and coherence analyses inside and outside watershed Areas

To compare different spectral dynamics of PSDs and coherence inside versus outside the watershed areas, the absolute values of interhemispheric PSD differences were summed across homotopic ROI pairs located either inside or outside watershed areas. These summed differences, |Σ(Δ_LR_ PSD)|, were computed separately for the two distinct frequency bands; 0.05 Hz and ∼0.1 Hz. The resulting values were then compared between the two groups (**Fig. 3C**). Similarly, coherence values between homotopic or heterotopic ROI pairs were calculated separately for inside and outside the watershed area. Fisher Z-transform was applied prior to averaging, and inverse Fisher Z-transform was applied after averaging coherence values across ROI pairs, separately for the two distinct frequency bands in each group (**Fig. 3D**, **S2 A**, and **S3 A**).

For network-specific analyses of FC inside and outside watershed areas, AICHA ROIs overlapping with the defined watershed territory were reordered according to the canonical rs-fMRI networks and subcortical regions. Network-specific coherence matrices were then averaged across the two distinct frequency bands (<0.05 Hz and ∼0.1 Hz), separately for regions inside and outside the watershed areas in each group. Fisher Z-transformation was applied prior to averaging coherence values across the networks, and inverse Z-transformation was performed after averaging. To visualize network-specific FC, circular diagrams were generated to represent the strength of coherence between the eight brain networks—comprising the seven canonical rs-fMRI networks and the subcortical network. Visualization was performed in R using ‘ggraph’ R library (package version 2.2.1). Detailed information on the AICHA ROIs overlapping with the distinct perfusion territories (i.e., inside vs. outside watershed areas) is provided in the Supplementary Information (**Table S2-3**).

## DATA PROCESSING FOR MULTIPARAMETRIC QUANTITATIVE MRI

### Estimation of CBV, CBF, CMRO_2_ and BOLD time lag in Gray Matter

Hemodynamic MRI parameter maps, i.e., relative CBV (from the DSC MRI), and CBF (from pCASL MRI), and relative OEF ≈ R2′/ CBV were calculated as described previously (Kaczmarz et al. 2021) (**Fig. S4**). Subsequently, relative CMRO_2_ maps were estimated using the following equation:

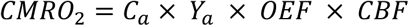

where *C_a_* is the oxygen carrying capacity of arterial blood (µmol O_2_ per 100 ml blood) and *Y_a_* is the hemoglobin oxygen saturation level of arterial blood. Following the work reported by An and colleagues (An et al. 2001), we assumed that *Y_a_* ≈ 1 and *C_a_* = 0.19 to simplify the estimation. To assess inter-hemispheric difference in BOLD time lag, lag time maps were generated from BOLD fMRI data using cross-correlation analysis in both groups. For each AICHA ROI, lag times were calculated as the time shift (ranging from -20 to +20 TRs) that maximized the cross-correlation coefficient with the fMRI signal from superior sagittal sinus. This analysis was implemented using the Python toolbox ‘rapiditide’, as described previously (Schneider et al. 2023). Detailed signal processing steps were provided in previous works (Schneider et al. 2022; Schneider et al. 2023).

### Inter-hemispheric differences in multiparametric quantitative MRI

To assess individual MRI parameter maps (CBV, CBF, CMRO_2_, and BOLD time lag), individual parameter maps were first parcellated using the AICHA atlas (Joliot et al. 2015). To compare differences between homotopic ROI pairs located inside versus outside the watershed areas, absolute inter-hemispheric differences were computed for each parameter. Specifically, for each subject, the absolute differences between left and right hemispheres were summed across homotopic ROI pairs using the following equation: |Σ(Δ_LR_ MRI Parameter)| = |Σ(MRI Parameter _Left_ - MRI Parameter _Right_|. This calculation was performed separately for ROIs inside and outside the watershed areas (**Fig. 3I-J**).

## STASTICAL ANALYSIS

To assess group differences in PSD and coherence, two-sample t-tests were performed to compare the mean values between healthy controls and patients with unilateral ICAS, considering p-values < 0.05 or < 0.01 statistically significant. For coherence analysis, Fisher-Z transform was applied to the coherence coefficient prior to the t-test to satisfy the model assumption (i.e., normal distribution). For comparison of inter-hemispheric differences in quantitative hemodynamic MRI parameters (CBV, CBF, CMRO_2_, and BOLD time lag), Wilcoxon rank-sum tests were used to evaluate differences between the two groups separately for inside and outside of watershed, using the same significant thresholds (p-values < 0.05 or < 0.01). In addition, effect sizes were calculated, i.e., Cohen’s d = (healthy control – patients with ICAS) / pooled SD.

## RESULTS

### Frequency-specific disruption of inter-hemispheric BOLD coherence at ∼0.1 Hz in patients with unilateral ICAS

We developed a multi-modal fMRI approach to assess whole-brain BOLD rs-fMRI signals and multi-parametric quantitative hemodynamic MRI maps to test our hypotheses in healthy controls and patients with asymptomatic unilateral ICAS (**Fig. 1A**). For inter-hemispheric ROI-based analysis, we employed an AICHA (Joliot et al. 2015), extracting fMRI time series from individual AICHA-defined ROIs (**Fig. 1B-C**). To test our first hypothesis that inter-hemispheric rs-fMRI coherence may be impaired in the ∼0.1 Hz frequency range (i.e., 0.08-0.1 Hz) (Mateo et al. 2017), we compared power spectral densities (PSDs) across hemispheres, as well as inter-hemispheric coherence, between healthy controls and patients with unilateral ICAS across the infra-slow oscillatory frequency band (<0.1 Hz). Using the AICHA atlas (Joliot et al. 2015) (**Fig. 1B**), we first analyzed the temporal oscillatory patterns and frequency responses of rs-fMRI signals between the right (ipsilateral to the stenosis in patients) and left hemispheres. In representative subjects from each group, we computed the ROI-average, Z-score normalized rs-fMRI time series for both hemispheres and compared the correlation and coherence values between them. As illustrated in **Fig. 1D and G**, the patient with ICAS exhibited weaker synchronization and less pronounced oscillatory patterns, resulting in lower correlation and coherence values, compared to the healthy control. To investigate alterations in inter-hemispheric FC in patients with ICAS, we analyzed Z-score normalized rs-fMRI time series for homotopic and heterotopic AICHA ROI pairs in the representative subjects shown in **Fig. 1G**. Based on correlation coefficients from the representative healthy control, we selected two AICHA ROI pairs, i.e., #1: Precuneus, #2: Lingual (**Fig. 1E-F**), which showed the highest homotopic correlation. We first examined homotopic rs-fMRI time series from the Precuneus ROI pair and compared correlation and coherence values between the two signals. As shown in **Fig. 1H**, the ICAS patient (right) exhibited synchronized but visibly weaker oscillatory temporal patterns compared to the healthy control (left). While the correlation coefficients were identical (control: 0.95; patient: 0.95), the coherence was notably reduced in the patient (control: 0.93; patient: 0.88), suggesting frequency-specific decoupling between the two hemispheres. Next, we assessed heterotopic rs-fMRI time series from the Precuneus and Lingual ROIs. As illustrated in **Fig. 1I**, the ICAS patient (right) showed markedly reduced synchronized and oscillatory temporal patterns, with both correlation and coherence values lower than those observed in the healthy control (left; correlation: control 0.72 vs. patient 0.53, coherence: control 0.51 vs. patient 0.40). These findings suggest that ICAS may differentially influence inter-hemispheric FC relying on the spatial relationship and spectral dynamics of the ROI pair.

To examine group-level frequency-specific effects, we computed PSDs separately for the right and left hemispheres. Both groups exhibited robust PSD peaks at 0.01-0.05 Hz (referred to as <0.05 Hz; **Fig. 1J**). However, in the ICAS group, a distinct alteration of PSD in the 0.08-0.1 Hz range (referred to as ∼0.1 Hz; **Fig. 1J**) was noted (see **Discussion**). Intriguingly, peaks at ∼0.02 Hz appeared only in ICAS patients, although this was not statistically significant (not shown), suggesting a potential spectral shift toward lower frequency (Drew et al. 2020; Kiviniemi et al. 2000). To statistically assess hemispheric difference, we calculated the absolute inter-hemispheric differences in PSDs (|Σ(Δ_LR_ PSD)|) for both frequency bands (<0.5 Hz and ∼0.1 Hz). Notably, a significant group difference emerged in the slow oscillatory frequency band, ∼0.1 Hz (p = 4.27 x 10^-2, Cohen’s d = −0.57), with ICAS patients showing stronger inter-hemispheric asymmetry compared to healthy controls (**Fig. 1K**), supporting a frequency-specific disruption of functional coherence in ICAS. We further conducted a group-level analysis of inter-hemispheric coherence across the infra-slow oscillatory frequency range (0.01-0.1 Hz, referred to as <0.1 Hz) in healthy controls and patients with unilateral ICAS. To compare coherence between groups, coherence values were Fisher Z-transformed to approximate a normal distribution, and Fisher Z-transformed coherence coefficients were computed between homotopic ROI pairs. As shown in **Fig. 1L**, group-averaged spectral coherence analysis revealed prominently reduced coherence values in the ICAS group, specifically in the 0.08-0.1 Hz range (∼0.1 Hz), while both groups exhibited similar coherence patterns in the ultra-slow oscillatory frequency band (0.01-0.05 Hz). Statistical analyses confirmed that homotopic coherence was significantly lower in patients with ICAS than in healthy controls in the 0.08-0.1 Hz band (p = 4.18 x 10^-2, Cohen’s d = 0.57, highlighted by a blue box in **Fig. 1M**). Similarly, heterotopic coherence analysis also showed a significant group difference at 0.08-0.1 Hz (p = 4.51 x 10^-2, Cohen’s d = 0.56, **Fig. 1N**).

In summary, supporting our first hypothesis, these findings indicate that the disruption of inter-hemispheric coherence in patients with ICAS is frequency-specific, with the most prominent differences occurring around ∼0.1 Hz. Moreover, the results clearly distinguish spectral coherence FC at ∼0.1 Hz between ICAS patients and healthy controls.

### Brain-wide mapping of inter-hemispheric coherence across canonical rs-fMRI networks in ultra-slow (<0.05 Hz) and slow (∼0.1 Hz) oscillatory frequency bands

Based on the inter-hemispheric PSD and coherence analysis, we examined the frequency-dependent coherence across large-scale networks in healthy controls and patients with unilateral ICAS. To define inter-hemispheric rs-fMRI networks, we employed seven canonical rs-fMRI networks proposed by Yeo and colleagues (Yeo et al. 2011). The seven rs-fMRI networks consisted of: DMN, SMN, DAN, VIN, SAN, LIN, and ECN. In addition, the SubCort was included in the analysis. Each AICHA ROI was assigned to one of these canonical rs-fMRI networks following the procedures described by Labache et al. (Labache et al. 2023) (**Fig. 2A** and **Table S1**). For network-level analysis of homotopic and heterotopic FC, we first visualized coherence matrices across the Yeo-defined networks for both groups at full infra-slow oscillatory frequency band (0.01-0.1 Hz). Interestingly, lower coherence values were observed in the ICAS group for heterotopic ROI pairs, particularly across VIN, SMN, and DAN (**Fig. 2B**). To examine frequency-specific patterns, we re-calculated inter-hemispheric coherence matrices at two distinct frequency bands: <0.05 Hz and ∼0.1 Hz. This analysis revealed notably diminished homotopic and heterotopic coherence across all networks in the ICAS group at ∼0.1 Hz, while coherence pattens at ultra-slow oscillatory frequency band (<0.05 Hz) remained largely comparable between the two groups (**Fig. 2C and E**).

To reduce noise and thus more clearly compare the coherence patterns across networks between the two groups, the ROI-based inter-hemispheric coherence matrices were averaged within each of the predefined networks (**Fig. 2D** and **2F**). At <0.05 Hz, both groups exhibited similar inter-hemispheric coherence patterns, with strong coherence observed along homotopic network pairs (diagonal) and across heterotopic connections involving the VIN, SMN, DAN, and SAN (upper four networks in the off-diagonal) (**Fig. 2D** and **S1 A**). No significant differences in inter-hemispheric coherence were observed between the two groups in this frequency band (**Fig. 2G**). In contrast, at ∼0.1 Hz, distinct coherence patterns emerged between the two groups and the ICAS group displayed globally reduced inter-hemispheric coherence, both in homotopic (diagonal) and heterotopic (off-diagonal) networks (**Fig. 2F and S1 B**). Statistically significant reductions in coherence were found in the VIN, SMN, DAN, and ECN in patients with ICAS (p = 4.57 x 10^-2, 3.81 x 10^-2, 1.83 x 10^-2, 3.34 x 10^-2; Cohen’s d = 0.56, 0.58, 0.67, 0.60, respectively), suggesting these cortical regions may share common cerebrovascular mechanisms (Kiviniemi et al. 2000) (**Fig. 2H**).

In summary, supporting our second hypothesis regarding spatially organized functional networks, these results demonstrate that frequency-specific disruptions in inter-hemispheric coherence at ∼0.1 Hz are consistently observed across the large-scale intrinsic brain networks. Moreover, the network-level coherence alterations at ∼0.1 Hz can clearly be distinguished from those at <0.05 Hz (ultra-slow oscillatory frequency band), highlighting the functional significance of frequency-dependent FC alteration in patients with ICAS.

### Regional vulnerability of inter-hemispheric BOLD coherence and vascular impairment inside watershed areas in patients with ICAS

To further explore regional effects on inter-hemispheric coherence, we investigated the impact of distinct perfusion conditions, i.e., inside vs. outside watershed areas, on frequency-dependent inter-hemispheric coherence in both healthy controls and patients with unilateral ICAS. As illustrated in **Fig. 3A**, we identified ROIs overlapping with pre-defined watershed areas using the AICHA atlas (Joliot et al. 2015) (n = 41 ROIs, details in the **Methods**). These ROIs were used to assess frequency-dependent inter-hemispheric coherence and to evaluate regional quantitative hemodynamic MRI parameters such CBV, CBF, CMRO_2_ and BOLD time lag (**Fig. 3A**). As shown in **Fig. 3B**, we quantified the distribution of the canonical rs-fMRI networks within the watershed areas: VIN: 54.3%, SMN: 4.8%, DAN: 15.8%, SAN: 0.0%, LIN: 22.2%, ECN: 38.9%, DMN: 10.0%, and SubCort: 13.6%. Notably, no ROI from SAN overlapped with the watershed area (**Table S2**).

To characterize spectral dynamics inside and outside watershed areas (**Table S2-3**), we computed group-averaged inter-hemispheric PSD differences (|Σ(Δ_LR_ PSD)|) at the two distinct frequency bands (ultra-slow; <0.5 Hz and slow; ∼0.1 Hz) for healthy controls and patients with ICAS. Notably, inside watershed areas, a significant group difference was observed only at ∼0.1 Hz (p = 4.84 x 10^-2, Cohen’s d = −0.56), whereas no significant differences were found at <0.05 Hz inside the watershed area, nor at either frequency band outside the watershed area (**Fig. 3C**). We also assessed group-averaged inter-hemispheric coherence in the same two frequency bands, again separately for regions inside and outside watershed areas. Consistent with the PSD results, significant group differences in both homotopic and heterotopic coherences were observed only at ∼0.1 Hz inside the watershed area (p = 1.40 x 10^-2 and 1.71 x 10^-2; Cohen’s d = 0.70 and 0.68, respectively). In contrast, coherence values at ∼0.1 Hz outside watershed areas, as well as at <0.05 Hz both inside and outside the watershed areas, did not differ significantly between the two groups (**Fig. 3D, S2, and S3**).

To further examine regional differences in inter-hemispheric coherence, we mapped coherence matrices across the canonical rs-fMRI networks separately for regions inside and outside watershed areas. For the two groups, network-averaged coherence matrices were computed at the two distinct frequency bands (<0.05 Hz and ∼0.1 Hz). As shown in **Fig. 3E**, inside the watershed areas, overall coherence values at ∼0.1 Hz were markedly diminished and spatially heterogeneous across the networks (with the exception of the SAN, which contained no ROI). The SubCort network exhibited low homotopic (diagonal) and heterotopic (off-diagonal) coherence values. These reductions are also visualized in the circular diagram, which illustrates pronounced decreases in inter-hemispheric coherence inside the watershed areas. Statistically, significantly lower coherence coefficients were observed in patients with ICAS across the seven cortical networks at ∼0.1 Hz (VIN, SMN, LIN, ECN, and DMN except DAN and SAN; SAN was not assigned, p = 2.76 x 10^-2, 4.50 x 10^-3, 3.80 x 10^-2, 4.11 x 10^-2, 1.84 x 10^-2; Cohen’s d = 0.62, 0.82, 0.59, 0.58, 0.67, respectively) (**Fig. 3G**), underscoring the impact of regional perfusion impairment on frequency-specific inter-hemispheric FC. In contrast, outside the watershed areas (**Fig. 3F**), coherence values at ∼0.1 Hz were only moderately reduced and were more homogeneous across heterotopic connections, while homotopic coherence remained relatively strong across most networks. The corresponding circular diagram confirms a lower degree of coherence reduction outside watershed areas compared to inside watershed areas. Only two networks, i.e., SMN and DAN, showed significantly lower inter-hemispheric coherence in patients with ICAS outside the watershed area (p = 4.92 x 10^-2 and 1.03 x 10^-2; Cohen’s d = 0.55 and 0.73, respectively) (**Fig. 3H**).

In addition, to investigate the impact of distinct perfusion territories (i.e., inside vs. outside watershed areas) on multi-parametric quantitative hemodynamic MRI parameters, we first estimated relative CBV, CBF, relative CMRO_2_, and BOLD time lag maps in healthy controls and patients with unilateral ICAS (**Fig. S4**). For each parameter, absolute inter-hemispheric differences were calculated by summing the difference between left and right hemispheres across homotopic ROI pairs: |Σ(Δ_LR_ CBV)|, |Σ(Δ_LR_ CBF)|, |Σ(Δ_LR_ CMRO_2_)|, and |Σ(Δ_LR_ BOLD lag)| (details in **Methods**). These metrics were then compared between the two groups, separately for inside and outside of watershed areas (**Fig. 3I-J**). Interestingly, the inter-hemispheric difference in relative CBV, |Σ(Δ_LR_ CBV)| showed a prominent and statistically significant group difference inside the watershed areas (p = 2.80 x 10^-3; Cohen’s d = −0.80), but not outside. In contrast, CBF differences were significant in both regions (inside: p = 1.71 x 10^-2; Cohen’s d = −0.51 and outside: p =1.55 x 10^-2; Cohen’s d = −0.70), showing diminished perfusion in the hemisphere ipsilateral to the stenosis (**Fig. S4 B**), while relative CMRO_2_ differences did not reach statistical significance in either region. This finding may indicate a generalized disruption of flow-metabolism coupling in patients, regardless of perfusion territory, consistent with previous findings (Gottler et al. 2019). For BOLD time lag, a significant inter-hemispheric difference was observed only outside the watershed areas (p = 3.28 x 10^-2; Cohen’s d = −0.61), while no significant difference was found inside the watershed areas, possibly reflecting asymmetric blood arrival times: the hemisphere ipsilateral to the stenosis show delayed perfusion outside the watershed territory, whereas both hemispheres exhibited more symmetric prolonged arrival times within the watershed areas (**Fig. S4 D**).

Taken together, in support of our third hypothesis regarding regions of distinct perfusion alterations, these findings demonstrate that aberrant inter-hemispheric rs-fMRI coherence at ∼0.1 Hz is localized specifically to vulnerable regions at the border of vascular territories, i.e., the watershed areas, that are particularly affected by hypoperfusion. Importantly, this ∼0.1 Hz-specific disruption in coherence appears to be independent of canonical rs-fMRI network organization. Furthermore, the results highlight the vulnerability of the watershed areas (Kaczmarz et al. 2021) in patients with unilateral ICAS, which are closely associated with a lateralized increase in CBV.

## DISCUSSION

Guided by our three hypotheses, i.e., frequency specificity, brain-wide effect, and regional vulnerability, this study demonstrates that frequency-specific alterations in inter-hemispheric PSD and coherence at ∼0.1 Hz distinguish patients with asymptomatic unilateral ICAS from healthy controls, despite largely preserved coherence at the ultra-slow oscillatory frequency band (<0.05 Hz). We further observed that inter-hemispheric coherence at ∼0.1 Hz is disrupted in a consistent way across canonical intrinsic networks and exhibits pronounced reduction in watershed areas, i.e., regions with particular perfusion vulnerability (Kaczmarz et al. 2021), where significant CBV lateralization is observed (p < 0.01). Together, these findings demonstrate a 0.1 Hz-specific impact of ICAS-related BOLD-FC alterations in watershed areas, where hemodynamic-vascular impairments are pronounced. This result is consistent with a model in which impaired vasomotor activity of ICAS – both widespread and more pronounced in watershed areas – may underlie the observed FC alterations.

Our multi-modal fMRI analysis reveals notably desynchronized time series, diminished coherence matrices, and lateralized PSDs at ∼0.1 Hz in patients with unilateral ICAS, accompanied by altered CBV inside watershed areas. Although these findings are consistent with hemodynamic and vasomotor-related impairments, the physiological interpretation of oscillations at ∼0.1 Hz requires caution due to methodological limitations. Given the relatively slow sampling rate of fMRI signals (TR = 1200 ms) and the absence of concurrent physiological signals recordings, the observed oscillations may reflect contributions from multiple sources, including vasomotor activity, systemic blood pressure fluctuations (i.e., Mayer waves), interhemispheric callosal projections, or other physiological confounders (Julien 2006; Mateo et al. 2017; Drew et al. 2020) (**Fig. 4**). First, vasomotor activity plays a central role in regulating arterial reservoir volumes and maintaining the homeostasis of flow and pressure in vascular networks (Nicoll and Webb 1955; Intaglietta 1990). However, since BOLD fMRI signals predominantly reflect venous rather than arterial dynamics (He et al. 2018), interpreting vasomotion influences on BOLD coherence requires careful consideration of vessel-specific contributions (Kim and Ogawa 2012). In particular, T2*-weighted EPI, which is commonly used for fMRI acquisition, is sensitive to large draining veins near superficial cortical layers (Choi et al. 2022; Choi et al. 2023; Choi et al. 2024; Tong and Frederick 2012). While arteries with smooth muscle cells generate vasomotor activity, capillaries (without muscle cells) lack such capability and only show passive responses due to homeostasis. Importantly, venules and veins also have smooth muscle cells (Boittin et al. 1999) and can regulate vasomotor activity, supporting venous return (Nicoll and Webb 1955; Aalkjær, Boedtkjer, and Matchkov 2011). Consistent with this view, prior studies have supported the existence of venous vasomotor activity (Jones 1852; Wiedeman 1957; Gotoh et al. 1982; Hundley et al. 1988; Wang et al. 2011). Nevertheless, future studies employing vessel-specific imaging approaches will be warranted to disentangle vessel-specific contributions to ∼0.1 Hz oscillation in rs-fMRI. Second, Mayer waves, which reflect systemic oscillation in arterial blood pressure, may overlap at the same frequency range (∼0.1 Hz) (Julien 2006; Elstad et al. 2011). Because these oscillations can coincide with BOLD fluctuations, changes in systemic and cerebrovascular hemodynamics may contribute to the observed rs-fMRI coherence to some extent (Hudetz, Roman, and Harder 1992; Julien 2006; Bumstead et al. 2017). Since blood pressure was not continuously monitored in this study, their potential contribution cannot be fully excluded. Third, the corpus callosum, a major fibers bundle connecting homotopic brain regions, is thought to mediate inter-hemispheric FC (Mohajerani et al. 2010; Magnuson et al. 2014; Johnston et al. 2008; Quigley et al. 2003). Reduced ∼0.1 Hz oscillation observed in acallosal mice has been interpreted as reflecting disrupted callosal projections (Mateo et al. 2017). However, the relative structural and functional contributions to ∼0.1 Hz oscillations remain to be clarified in future studies (Schmitzer et al. 2024; Avena-Koenigsberger, Misic, and Sporns 2017). Fourth, other physiological confounders should also be taken into account. Cardiac pulsations (∼1 Hz) could introduce aliasing artifacts near ∼0.1 Hz, particularly given the Nyquist sampling limit of rs-fMRI signals (∼0.4 Hz) (Chang, Cunningham, and Glover 2009; Birn 2012; Caballero-Gaudes and Reynolds 2017). Future studies incorporating concurrent physiological recording or faster fMRI acquisition methods will be necessary to fully rule out physiological noise contributions. In addition, the temporal bandpass filtering during preprocessing (0.01-0.1 Hz) may modulate the spectral profile near the upper boundary of this range. As shown by He et al (He et al. 2010) and Choi et al (Choi et al. 2023), PSD analyses showed that spectral responses exhibit greater variability around ∼0.1 Hz, following 1/f power regime (Biswal et al. 1995; He et al. 2018; Choi et al. 2022; Choi et al. 2023; He 2014). Spectral peaks can also be influenced by frequency resolution and smoothing procedure applied during preprocessing (Muthuswamy and Thakor 1998; van Vugt, Sederberg, and Kahana 2007; Kleinfeld and Mitra 2014). Furthermore, behavioral state fluctuation such as sleep or fidgeting may have contributed to variability at ∼0.1 Hz (Lei et al. 2015; Drew et al. 2020; Helakari et al. 2022; Fultz et al. 2019; Hauglund et al. 2025), especially in this elderly cohort during relatively long scanning sessions.

**Figure 4.**
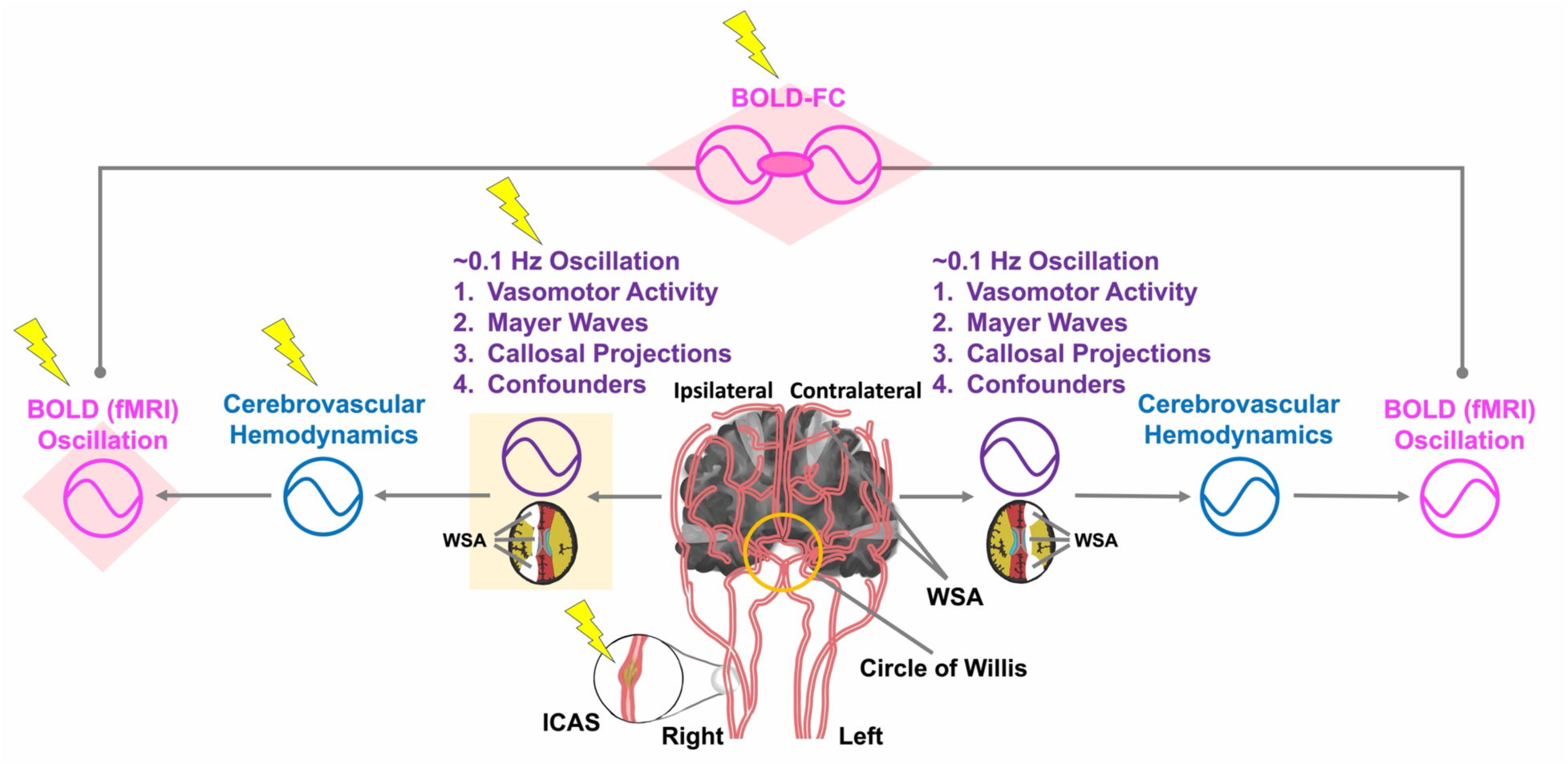
Interpretational cascade model of unilateral ICAS-driven cerebrovascular hemodynamics and its impact on BOLD functional connectivity (FC) inspired by the seminal work of Mateo et al. (Mateo et al. 2017). Unilateral internal carotid artery stenosis (ICAS, lightning symbol) may lead to impaired ∼0.1 Hz oscillations transmitted through shared vascular networks, particularly via the Circle of Willis. Disrupted ∼0.1 Hz oscillations, arising from vasomotor activity, Mayer waves, callosal projections, and other physiological confounders, in the hemisphere ipsilateral to ICAS alter downstream cerebrovascular hemodynamics (e.g., cerebral blood flow, cerebral blood volume, and cerebral metabolic rate of oxygen), especially in vulnerable watershed areas (WSAs). These hemodynamic alterations result in disrupted frequency-specific BOLD oscillations and FC between two hemispheres, as measured by reduced inter-hemispheric coherence at ∼0.1 Hz.

Although vasomotor oscillations span a broad frequency range (0.01 to 0.3 Hz, centered near 0.1 Hz) (Aalkjær, Boedtkjer, and Matchkov 2011; Drew et al. 2020), respiratory aliasing artifacts often confound fMRI signals in the 0.15-0.3 Hz range in human fMRI (Glover, Li, and Ress 2000; Birn, Murphy, and Bandettini 2008; Birn 2012; Caballero-Gaudes and Reynolds 2017), especially without concurrent respiratory recording or physiological noise correction. To minimize contamination from aliased respiratory signals and maximize physiological interpretability, we therefore focused on slow fluctuations within the commonly used resting-state frequency band of 0.01-0.1 Hz (Fox and Raichle 2007). Within this range, we define the ∼0.1 Hz band as the 0.08-0.1 Hz to probe vasomotor-related fluctuations, as spectral peaks in this range have been consistently observed in prior human studies using fMRI (Mitra et al. 1997), near infrared spectroscopy (Obrig et al. 2000), and multi-spectral optical intrinsic signal imaging (Rayshubskiy et al. 2014). To mor directly address causality, future studies in ICAS should compare ∼0.1 Hz-specific coherence before and after post-revascularization (Cheng et al. 2012) or incorporate concurrent transcranial Doppler measurements to directly assess vasomotor activity alongside the present multi-modal fMRI framework. Complementary animal models with experimentally induced unilateral stenosis (Ishikawa et al. 2023) (see **Fig. S6**), combined with concurrent optical recordings, could further provide direct evidence for the relationship between cerebrovascular impairment and inter-hemispheric coherence disruption.

In contrast to this clear FC disruption at ∼0.1 Hz, ultra-slow oscillations (<0.05 Hz) remained stable in patients with ICAS (**Fig. 1L**, **2C-D**, and **2G**). This stable ultra-slow oscillations, together with preserved CMRO_2_ (**Fig. 3I-J**), suggest preserved neuronal activity (He et al. 2018) and global arousal brain state via neuromodulators, such as adrenergic and cholinergic neurons (Raut et al. 2021; Reimer et al. 2016). This pattern followed a spatial gradient from strong to weak coherence, spanning from unimodal (VIN and SMN), multi-modal (DAN and SAN), to transmodal (ECN and DMN) networks, reflecting known functional hierarchies and principal gradient axes (Raut et al. 2021; Margulies et al. 2016). Moreover, the preserved ultra-slow oscillations align with previous laminar fMRI findings (Choi et al. 2022; Choi et al. 2023), and differences in ultra-slow coherence between cortical and subcortical structures may reflect distinct neurovascular coupling mechanisms (**Fig. 2D**). However, since we did not directly measure neuronal activity, the interpretation of preserved CMRO_2_ requires caution. In this study, CMRO_2_ was estimated from relative OEF (≈ R2′/ CBV) and CBF, as described in the **Methods** (Kaczmarz et al. 2021) (**Fig. S4**). Thus, preserved CMRO_2_ may reflect opposing changes of OEF and CBF in the hemisphere ipsilateral to ICAS. Concurrent neuronal recording combined with human fMRI would help clarify the relationship between neuronal activity, ultra-slow oscillations (<0.05 Hz), and slow oscillations (∼0.1 Hz) as has been shown in animal studies (Mateo et al. 2017; He et al. 2018; Broggini et al. 2024).

Watershed regions are of particular clinical relevance in ICAS patients (Torvik 1984; Kaczmarz et al. 2018; Kaczmarz et al. 2021). Inside watershed areas, coherence matrices at ∼0.1 Hz showed pronounced spatial heterogeneity across networks, consistent with the more severe hemodynamic impairments reported in prior work (Kaczmarz et al. 2018; Kaczmarz et al. 2021) (**Fig. 3E**). This heterogeneity likely reflects an interplay between compensation mechanisms (e.g., collateralization and autoregulation) (Zarrinkoob et al. 2019; Reinhard et al. 2003) and cerebrovascular impairment (Kaczmarz et al. 2018; Kaczmarz et al. 2021). In contrast, coherence at ∼0.1 Hz outside watershed areas was spatially homogeneous and relatively preserved (**Fig. 3F**), although diminished coherence at ∼0.1 Hz was observed in SMN and DAN outside watershed areas, potentially due to age-related limitations in compensatory capacity in ICAS patients (**Fig. 3H**). These observations suggest that ∼0.1 Hz-related coherence could serve as a sensitive marker of regional vulnerability to vascular impairment in ICAS patients, highlighting the increased sensitivity inside watershed areas.

A major methodological strength of this study is the integration of frequency-specific rs-fMRI analysis with multi-parametric hemodynamic MRI, enabling the identification of ∼0.1 Hz-specific oscillatory disruptions in functional dynamics and region-specific vascular vulnerabilities, particularly inside watershed areas. By examining PSD and coherence at two distinct frequency bands, i.e., ultra-slow (<0.05 Hz) and slow (∼0.1 Hz), and linking these metrics to quantitative maps of CBV, CBF, CMRO_2_, and BOLD time lag, we observed that inter-hemispheric desynchronization at ∼0.1 Hz is specifically associated with ipsilaterally increased CBV, as a potential factor related to impaired hemodynamics due to chronic vasodilation. In contrast, conventional correlation-based FC measures did not reveal significant group differences between healthy controls and ICAS patients (Schneider et al. 2022; Schneider et al. 2023) (**Fig. S5**), likely due to their limited sensitivity to frequency-specific impacts on FC. Together, our multi-modal fMRI framework extends beyond conventional correlation-based approaches by enabling the investigation of brain-wide, frequency-specific functional dynamics and vascular contributions to rs-fMRI signals in ICAS patients. In the future, coherence-based fMRI approaches may offer valuable insights for detecting subtle vascular impairment and functional desynchronization in patients with neurological and neurovascular disorders, including but not limited to ICAS.

In conclusion, our findings suggest that inter-hemispheric BOLD coherence at ∼0.1 Hz may serve as a frequency-sensitive and spatially specific indicator of cerebrovascular impairment in patients with asymptomatic unilateral ICAS. By linking frequency-specific functional coherence to regional hemodynamic alterations, this work helps to bridge the gap between vascular pathology and resting-state functional disruption, and highlight the potential of coherence-based spectral analysis as a distinct, non-invasive index not only for ICAS but also for other clinical conditions associated with vascular impairment, including stroke, small vessel disease, dementia, and Alzheimer’s Disease (van Veluw et al. 2020; Zhang et al. 2024; Wardlaw, Smith, and Dichgans 2019; Holstein-Ronsbo et al. 2023; Di Marco et al. 2015).

### Data availability

Availability of human subject data is restricted due to data privacy regulations. Data sharing requires specific ethics votes and data transfer agreements. Processed non-image data generated during this study are available from the corresponding author upon reasonable request.

### Code availability

The related image processing codes are available from the corresponding author upon reasonable request.

## Acknowledgements

We thank Jacob Duckworth, Drs. David Kleinfeld, Filip Sobczak, Loïc Labache, and Afra Wohlschläger for their helpful comments and discussions. We also thank Yehwan (Issac) Choi for his schematic illustrations. This work was supported the Deutsche Forschungsgemeinschaft (DFG, German Research Foundation) – project number 395030489 and ev. Studienwerk Villigst (personal grant to Gabriel Hoffmann).

## Author contributions

C.P. and C.S. designed and supervised the research, S.K. and C.P. performed experiments and acquired data, S.C. analyzed data, G.H., and S.S. provided technical support, S.C., X.Y., C.S., and C.P. wrote the manuscript.

## Competing interests

S.K. is an employee of Philips GmbH Market DACH, Hamburg, Germany. X.Y. is a co-founder of MRIBOT LLC. The remaining authors declare no conflict of interest.

## Notes

### Summary of Updates

We revised the abstract, introduction, and discussion.

