## Supplementary Information for "Resting-state fMRI coherence is selectively diminished around 0.1 Hz, particularly inside watershed areas, in patients with unilateral carotid artery stenosis"

**Title:**

**Supplementary Table:** 4

**Supplementary Figure:** 6

***Lead corresponding author:**
Dr. Sangcheon Choi


Address: Ismaninger Str. 22, 81675 Munich, Germany

**Table S1.** Assignment of Individual AICHA ROIs (N = 192) to seven canonical resting-state networks. ROIs are symmetrically distributed across the right and left hemispheres.

| Networks | Assigned AICHA ROIs |
|---|---|
| Visual Network (VIN) | 'G_Occipital_Pole-1', 'G_Occipital_Lat-1', 'G_Occipital_Lat-2', 'G_Occipital_Lat-3', 'G_Occipital_Lat-4', 'G_Occipital_Lat-5', 'G_Occipital_Sup-1', 'G_Occipital_Sup-2', 'G_Occipital_Mid-1', 'G_Occipital_Mid-2', 'G_Occipital_Inf-1', 'G_Occipital_Inf-2', 'G_Temporal_Inf-5', 'S_Parietooccipital-1', 'S_Parietooccipital-4', 'S_Parietooccipital-5', 'S_Parietooccipital-6', 'G_Cuneus-1', 'G_Cuneus-2', 'G_Calcarine-1', 'G_Calcarine-2', 'G_Calcarine-3', 'G_Lingual-1', 'G_Lingual-2', 'G_Lingual-3', 'G_Lingual-4', 'G_Lingual-5', 'G_Lingual-6', 'G_ParaHippocampal-2', 'G_ParaHippocampal-4', 'G_ParaHippocampal-5', 'G_Fusiform-3', 'G_Fusiform-5', 'G_Fusiform-6', 'G_Fusiform-7' |
| Somato-Motor Network (SMN) | 'S_Precentral-3', 'S_Precentral-6', 'S_Rolando-1', 'S_Rolando-2', 'S_Rolando-3', 'S_Rolando-4', 'S_Postcentral-1', 'S_Postcentral-2', 'G_Parietal_Sup-1','G_Parietal_Sup-2', 'G_Insula-posterior-1', 'G_Rolandic_Oper-2', 'G_Temporal_Sup-1', 'G_Temporal_Sup-2', 'G_Temporal_Sup-3', 'S_Cingulate-4', 'S_Cingulate-7', 'G_Paracentral_Lobule-1', 'G_Paracentral_Lobule-2', 'G_Paracentral_Lobule-3', 'G_Paracentral_Lobule-4' |
| Dorsal Attention Network (DAN) | 'G_Frontal_Sup-3', 'S_Sup_Frontal-6', 'S_Precentral-1', 'S_Precentral-2', 'S_Precentral-4', 'S_Precentral-5', 'S_Postcentral-3', 'G_Parietal_Sup-3', 'G_Parietal_Sup-4', 'G_Parietal_Sup-5', 'S_Intraparietal-3', 'S_Intraoccipital-1', 'G_Occipital_Mid-3', 'G_Temporal_Mid-4', 'G_Temporal_Inf-3', 'G_Temporal_Inf-4', 'G_Precuneus-6', 'G_Precuneus-9', 'G_Fusiform-4' |
| Salience Network (SAN) | 'S_Sup_Frontal-3', 'G_Supramarginal-1', 'G_SupraMarginal-2', 'G_Supramarginal-3', 'G_Supramarginal-4', 'G_SupraMarginal-5', 'G_Insula-anterior-3', 'G_Insula-anterior-4', 'G_Insula-anterior-5', 'G_Rolandic_Oper-1', 'S_Sup_Temporal-4', 'G_Temporal_Pole_Sup-2', 'S_Cingulate-1', 'S_Cingulate-2', 'S_Cingulate-3','S_Cingulate-5', 'S_Cingulate-6', 'G_Cingulum_Mid-1', 'G_Precuneus-5' |
| Limbic Network (LIN) | 'S_Sup_Frontal-1', 'G_Frontal_Sup_Orb-1', 'G_Frontal_Inf_Orb-2', 'S_Orbital-1', 'S_Olfactory-1', 'G_Insula-anterior-1', 'G_Temporal_Inf-1', 'G_Temporal_Pole_Sup-1', 'G_Temporal_Pole_Mid-1', 'G_Temporal_Pole_Mid-2', 'G_Temporal_Pole_Mid-3', 'G_Frontal_Med_Orb-1', 'G_subcallosal-1', 'G_Hippocampus-1', 'G_ParaHippocampal-3', 'G_Fusiform-1', 'G_Fusiform-2', 'N_Caudate-3' |
| Executive Control Network (ECN) | 'S_Sup_Frontal-2', 'G_Frontal_Mid-1', 'G_Frontal_Mid-2', 'G_Frontal_Mid-3', 'G_Frontal_Mid-5', 'S_Inf_Frontal-1', 'S_Inf_Frontal-2', 'G_Frontal_Mid_Orb-1', 'G_Frontal_Mid_Orb-2', 'G_SupraMarginal-6', 'G_Parietal_Inf-1', 'S_Intraparietal-1', 'S_Intraparietal-2', 'G_Frontal_Sup_Medial-3', 'G_Supp_Motor_Area-1', 'G_Cingulum_Mid-2', 'G_Cingulum_Post-1', 'S_Parietooccipital-3' |
| Default Mode Network (DMN) | 'G_Frontal_Sup-1', 'G_Frontal_Sup-2', 'S_Sup_Frontal-4', 'S_Sup_Frontal-5', 'G_Frontal_Mid-4', 'G_Frontal_Inf_Tri-1', 'G_Frontal_Inf_Orb-1', 'S_Orbital-2', 'G_SupraMarginal-7', 'G_Angular-1', 'G_Angular-2', 'G_Angular-3', 'G_Occipital_Mid-4', 'G_Insula-anterior-2', 'G_Temporal_Sup-4', 'S_Sup_Temporal-1', 'S_Sup_Temporal-2', 'S_Sup_Temporal-3', 'S_Sup_Temporal-5', 'G_Temporal_Mid-1', 'G_Temporal_Mid-2', 'G_Temporal_Mid-3', 'G_Temporal_Inf-2', 'G_Frontal_Sup_Medial-1', 'G_Frontal_Sup_Medial-2', 'S_Anterior_Rostral-1', 'G_Frontal_Med_Orb-2', 'G_Supp_Motor_Area-2', 'G_Supp_Motor_Area-3', 'G_Cingulum_Ant-1', 'G_Cingulum_Ant-2', 'G_Cingulum_Mid-3', 'G_Cingulum_Post-2', 'G_Precuneus-1', 'G_Precuneus-2', 'G_Precuneus-3', 'G_Precuneus-4', 'G_Precuneus-7', 'G_Precuneus-8', 'S_Parietooccipital-2' |
| Sub-Cortical Region (SubCort) | 'G_Cingulum_Post-3', 'G_Hippocampus-2', 'G_ParaHippocampal-1', 'N_Amygdala-1', 'N_Caudate-1', 'N_Caudate-2', 'N_Caudate-4', 'N_Caudate-5', 'N_Caudate-6', 'N_Caudate-7', 'N_Pallidum-1', 'N_Putamen-2', 'N_Putamen-3', 'N_Thalamus-1', 'N_Thalamus-2', 'N_Thalamus-3', 'N_Thalamus-4', 'N_Thalamus-5', 'N_Thalamus-6', 'N_Thalamus-7', 'N_Thalamus-8', 'N_Thalamus-9' |

**Table S2**. Assignment of Individual AICHA ROIs (N = 41) to seven canonical resting-state networks located inside watershed areas. ROIs are symmetrically distributed across the right and left hemispheres.

| Networks | Assigned AICHA ROIs inside watershed areas |
|---|---|
| Visual Network (VIN) | 'G_Occipital_Pole-1', 'G_Occipital_Lat-2', 'G_Occipital_Lat-3', 'G_Occipital_Lat-4', 'G_Occipital_Lat-5', 'G_Occipital_Sup-1', 'G_Occipital_Sup-2', 'G_Occipital_Mid-1', 'G_Occipital_Mid-2', 'G_Occipital_Inf-1', 'G_Occipital_Inf-2', 'G_Temporal_Inf-5', 'S_Parietooccipital-1', 'S_Parietooccipital-4', 'S_Parietooccipital-6', 'G_Cuneus-1', 'G_Cuneus-2', 'G_Calcarine-1', 'G_Fusiform-7' |
| Somato-Motor Network (SMN) | 'S_Precentral-3' |
| Dorsal Attention Network (DAN) | 'S_Intraoccipital-1', 'G_Occipital_Mid-3', 'G_Precuneus-9' |
| Salience Network (SAN) | N/A |
| Limbic Network (LIN) | 'G_Insula-anterior-1', 'G_Temporal_Inf-1', 'G_Temporal_Pole_Sup-1', 'N_Caudate-3' |
| Executive Control Network (ECN) | 'G_Frontal_Mid-2', 'G_Frontal_Mid-3', 'G_Frontal_Mid-5', 'S_Inf_Frontal-1', 'S_Inf_Frontal-2', 'G_Parietal_Inf-1', 'S_Parietooccipital-3' |
| Default Mode Network (DMN) | 'G_Frontal_Mid-4', 'G_Frontal_Inf_Tri-1', 'G_Occipital_Mid-4', 'G_Precuneus-8' |
| Sub-Cortical Region (SubCort) | 'N_Amygdala-1', 'N_Caudate-7', 'N_Pallidum-1' |

**Table S3**. Assignment of Individual AICHA ROIs (N = 151) to seven canonical resting-state networks located outside watershed areas. ROIs are symmetrically distributed across the right and left hemispheres.

| Networks | Assigned AICHA ROIs outside watershed areas |
|---|---|
| Visual Network (VIN) | 'G_Occipital_Lat-1', 'S_Parietooccipital-5', 'G_Calcarine-2', 'G_Calcarine-3', 'G_Lingual-1', 'G_Lingual-2', 'G_Lingual-3', 'G_Lingual-4', 'G_Lingual-5', 'G_Lingual-6', 'G_ParaHippocampal-2', 'G_ParaHippocampal-4', 'G_ParaHippocampal-5', 'G_Fusiform-3', 'G_Fusiform-5', 'G_Fusiform-6' |
| Somato-Motor Network (SMN) | 'S_Precentral-6', 'S_Rolando-1', 'S_Rolando-2', 'S_Rolando-3', 'S_Rolando-4', 'S_Postcentral-1', 'S_Postcentral-2', 'G_Parietal_Sup-1', 'G_Parietal_Sup-2', 'G_Insula-posterior-1', 'G_Rolandic_Oper-2', 'G_Temporal_Sup-1', 'G_Temporal_Sup-2', 'G_Temporal_Sup-3', 'S_Cingulate-4', 'S_Cingulate-7', 'G_Paracentral_Lobule-1', 'G_Paracentral_Lobule-2', 'G_Paracentral_Lobule-3', 'G_Paracentral_Lobule-4' |
| Dorsal Attention Network (DAN) | 'G_Frontal_Sup-3', 'S_Sup_Frontal-6', 'S_Precentral-1', 'S_Precentral-2', 'S_Precentral-4', 'S_Precentral-5', 'S_Postcentral-3', 'G_Parietal_Sup-3', 'G_Parietal_Sup-4', 'G_Parietal_Sup-5', 'S_Intraparietal-3', 'G_Temporal_Mid-4', 'G_Temporal_Inf-3', 'G_Temporal_Inf-4', 'G_Precuneus-6', 'G_Fusiform-4' |
| Salience Network (SAN) | 'S_Sup_Frontal-3', 'G_Supramarginal-1', 'G_SupraMarginal-2', 'G_Supramarginal-3', 'G_Supramarginal-4', 'G_SupraMarginal-5', 'G_Insula-anterior-3', 'G_Insula-anterior-4', 'G_Insula-anterior-5', 'G_Rolandic_Oper-1', 'S_Sup_Temporal-4', 'G_Temporal_Pole_Sup-2', 'S_Cingulate-1', 'S_Cingulate-2', 'S_Cingulate-3', 'S_Cingulate-5', 'S_Cingulate-6', 'G_Cingulum_Mid-1', 'G_Precuneus-5' |
| Limbic Network (LIN) | 'S_Sup_Frontal-1', 'G_Frontal_Sup_Orb-1', 'G_Frontal_Inf_Orb-2', 'S_Orbital-1', 'S_Olfactory-1', 'G_Temporal_Pole_Mid-1', 'G_Temporal_Pole_Mid-2', 'G_Temporal_Pole_Mid-3', 'G_Frontal_Med_Orb-1', 'G_subcallosal-1', 'G_Hippocampus-1', 'G_ParaHippocampal-3', 'G_Fusiform-1', 'G_Fusiform-2' |
| Executive Control Network (ECN) | 'S_Sup_Frontal-2', 'G_Frontal_Mid-1', 'G_Frontal_Mid_Orb-1', 'G_Frontal_Mid_Orb-2', 'G_SupraMarginal-6', 'S_Intraparietal-1', 'S_Intraparietal-2', 'G_Frontal_Sup_Medial-3', 'G_Supp_Motor_Area-1', 'G_Cingulum_Mid-2', 'G_Cingulum_Post-1' |
| Default Mode Network (DMN) | 'G_Frontal_Sup-1', 'G_Frontal_Sup-2', 'S_Sup_Frontal-4', 'S_Sup_Frontal-5', 'G_Frontal_Inf_Orb-1', 'S_Orbital-2', 'G_SupraMarginal-7', 'G_Angular-1', 'G_Angular-2', 'G_Angular-3', 'G_Insula-anterior-2', 'G_Temporal_Sup-4', 'S_Sup_Temporal-1', 'S_Sup_Temporal-2', 'S_Sup_Temporal-3', 'S_Sup_Temporal-5', 'G_Temporal_Mid-1', 'G_Temporal_Mid-2', 'G_Temporal_Mid-3', 'G_Temporal_Inf-2', 'G_Frontal_Sup_Medial-1', 'G_Frontal_Sup_Medial-2', 'S_Anterior_Rostral-1', 'G_Frontal_Med_Orb-2', 'G_Supp_Motor_Area-2', 'G_Supp_Motor_Area-3', 'G_Cingulum_Ant-1', 'G_Cingulum_Ant-2', 'G_Cingulum_Mid-3', 'G_Cingulum_Post-2', 'G_Precuneus-1', 'G_Precuneus-2', 'G_Precuneus-3', 'G_Precuneus-4', 'G_Precuneus-7', 'S_Parietooccipital-2' |
| Sub-Cortical Region (SubCort) | 'G_Cingulum_Post-3', 'G_Hippocampus-2', 'G_ParaHippocampal-1', 'N_Caudate-1', 'N_Caudate-2', 'N_Caudate-4', 'N_Caudate-5', 'N_Caudate-6', 'N_Putamen-2', 'N_Putamen-3', 'N_Thalamus-1', 'N_Thalamus-2', 'N_Thalamus-3', 'N_Thalamus-4', 'N_Thalamus-5', 'N_Thalamus-6', 'N_Thalamus-7', 'N_Thalamus-8', 'N_Thalamus-9' |

**Table S4**. Demographic information.

| | Control (N=26) | ICAS (N=27) | P-value |
|---|---|---|---|
| **Age (years): mean ± SD** | 70.3 ± 4.8 | 70.3 ± 7.2 | 0.994 |
| **Sex – female: N** | 15 (57.7%) | 11 (40.7%) | 0.217 |
| **Side of Stenosis – Right: N** | 0 (0.0%) | 16 (59.3%) | **< 0.001*** |
| **NASCET: mean ± SD** | N/A | 81.0 ± 9.8 % | |
| **BMI: : mean ± SD** | 27.0 ± 4.2 | 26.0 ± 4.7 | 0.414 |
| **Smoking (packs/year): mean ± SD** | 6.8 ± 12.3 | 12.9 ± 18.4 | 0.165 |
| **Fazekas: mean ± SD** | 0.9 ± 0.9 | 1.5 ± 0.8 | **0.014** |
| **PAOD: N** | 3 (11.5%) | 8 (29.6%) | 0.104 |
| **CHD: N** | 2 (7.7%) | 9 (34.6%) | **0.017** |
| **Hypertension** | 15 (57.7%) | 21 (77.8%) | 0.117 |
| **BP (mmHg)** | | | |
| **- systolic: mean ± SD** | 142.1 ± 18.9 | 153.6 ± 25.3 | 0.072 |
| **- diastolic: mean ± SD** | 84.9 ± 7.0 | 85.5 ± 10.3 | 0.797 |
| **Diabetes: N** | 2 (7.7%) | 6 (23.1%) | 0.124 |
| **Statins: N** | 5 (19.2%) | 19 (70.4%) | **< 0.001** |
| **Antiplatelets: N** | 5 (19.2%) | 24 (88.9%) | **< 0.001** |
| **Antihypertensives: N** | 11 (42.3%) | 18 (66.7%) | 0.075 |
| **Antidiabetics: N** | 2 (7.7%) | 3 (11.1%) | 0.670 |
| **MMSE: mean ± SD** | 28.6 ± 1.4 | 28.3 ± 2.4 | 0.607 |
| **TMT-A: mean ± SD** | 42.3 ± 13.2 | 49.7 ± 23.8 | 0.175 |
| **TMT-B: mean ± SD** | 103.2 ± 40.1 | 139.0 ± 65.0 | **0.023** |

| | | | |
|---|---|---|---|
| **LBT (absolute deviation from center) : mean ± SD** | 3.1 ± 1.8 | 2.4 ± 2.2 | 0.203 |
| **BDI: mean ± SD** | 8.5 ± 5.2 | 8.7 ± 8.1 | 0.921 |
| **STAI: mean ± SD** | 35.0 ± 10.3 | 37.9 ± 10.4 | 0.323 |

Note: Variables are presented as mean ± standard deviation (SD) or as absolute number (N) with corresponding percentage (%). Group comparisons for age, mean pack-years, blood pressure (BP), body mass index (BMI), Mini-Mental State Examination (MMSE), Trail Making Test A/B (TMT-A/B), Beck Depression Inventory (BDI), State-Trait Anxiety Inventory (STAI), and Line Bisection Test (LBT) were performed using two-sample t-tests. Chi-squared tests were applied to all other categorical variables. Statistically significant group differences ($p < 0.05$) are indicated in **bold**. Abbreviations: BMI, body mass index; PAOD, peripheral artery occlusive disease; CHD, coronary heart disease; BP, blood pressure; MMSE, Mini-Mental State Examination; TMT-A/B, Trail Making Test A/B; LBT, Line Bisection Test; BDI, Beck Depression Inventory; STAI, State-Trait Anxiety Inventory.

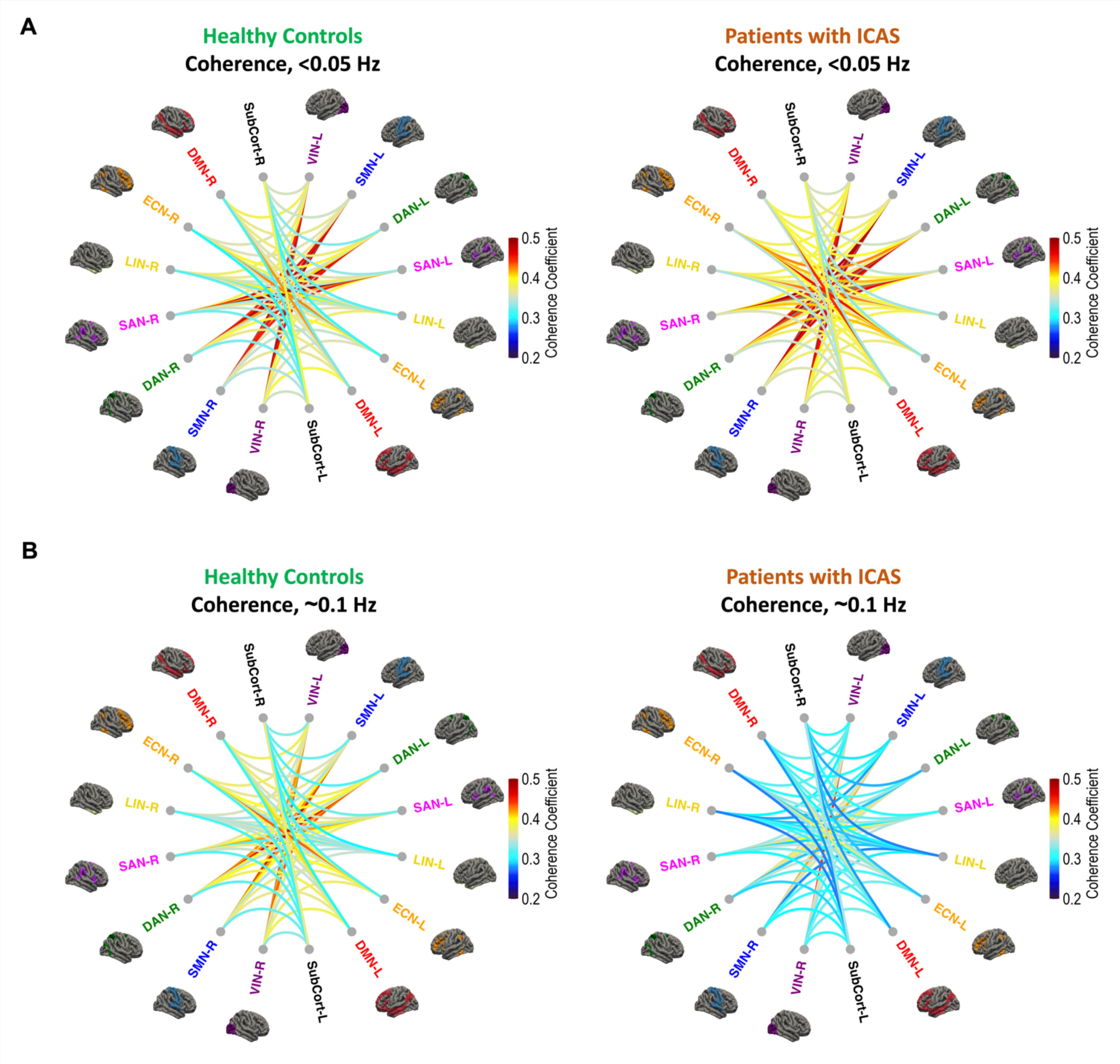


**Figure S1.** Circular visualizations corresponding to network-wise inter-hemispheric coherence matrix in healthy controls and patients with ICAS, corresponding to **Fig. 3D** and **F**. **A.** Inter-hemispheric coherence at ultra-slow frequency band (<0.05 Hz). **B.** Inter-hemispheric coherence at vasomotor frequency band (~0.1 Hz).

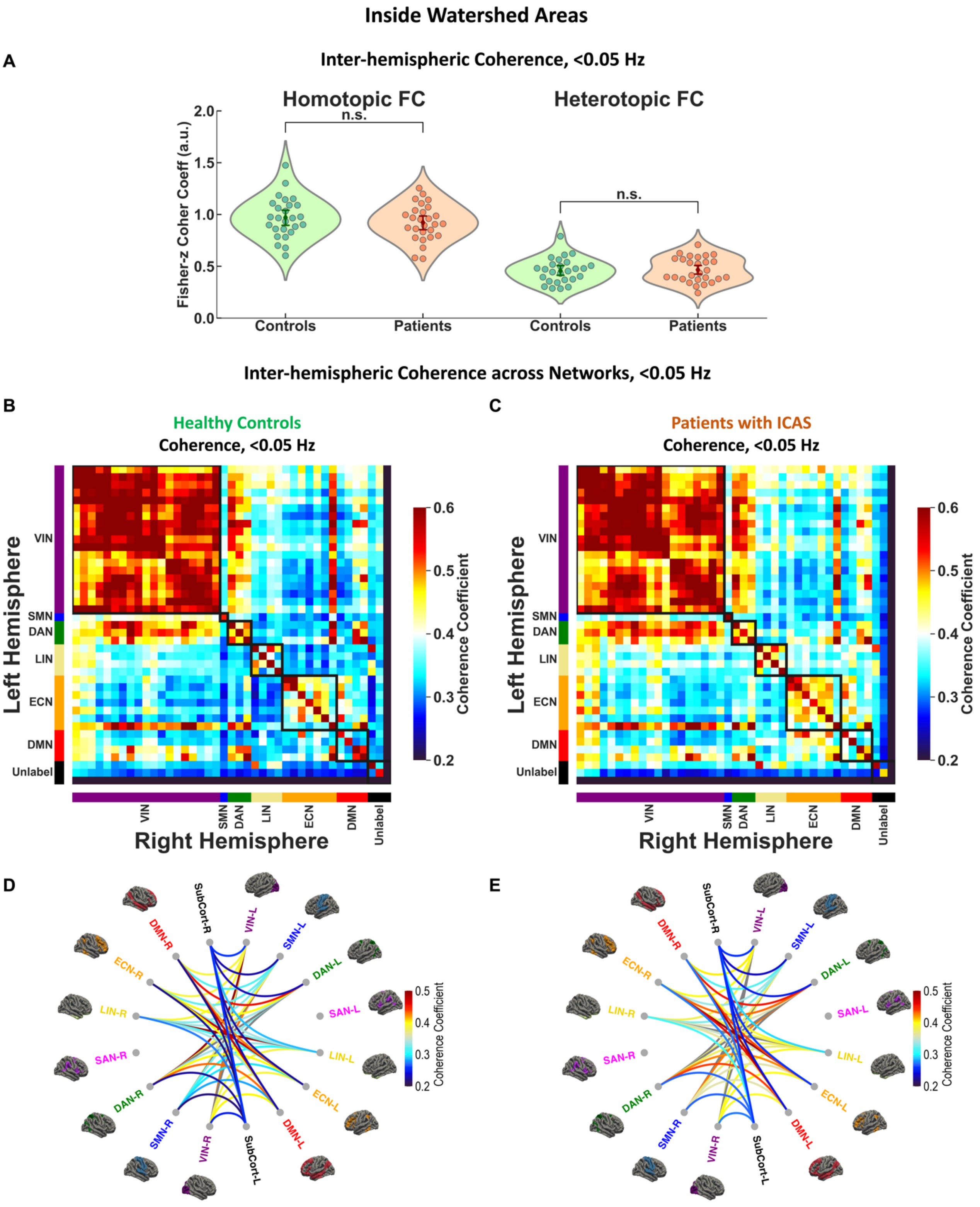

Inside Watershed Areas
A
Inter-hemispheric Coherence, <0.05 Hz
Homotopic FC
Heterotopic FC
n.s.
n.s.
Fisher-z Coher Coeff (a.u.)
Controls
Patients
Controls
Patients
Inter-hemispheric Coherence across Networks, <0.05 Hz
B
Healthy Controls
Coherence, <0.05 Hz
C
Patients with ICAS
Coherence, <0.05 Hz
Left Hemisphere
Right Hemisphere
VIN
SMN
DAN
LIN
ECN
DMN
Unlabel
Coherence Coefficient
D
E
SubCort-R
VIN-L
SMN-L
DAN-L
SAN-L
LIN-L
ECN-L
DMN-L
SubCort-L
VIN-R
SMN-R
DAN-R
SAN-R
LIN-R
ECN-R
DMN-R

**Figure S2.** Group-level comparisons of inter-hemispheric coherence inside watershed areas at <0.05 Hz. **A.** Inter-hemispheric coherence values for homotopic and heterotopic FCs in healthy controls (green) and patients with ICAS (orange). **B-C.** Network-level inter-hemispheric coherence matrices in healthy controls (**B**) and patients with ICAS (**C**). **D-E.** Corresponding circular visualizations for healthy controls (**D**) and patients with ICAS (**E**). Note that the SAN is not assigned in this analysis.

### Outside Watershed Areas

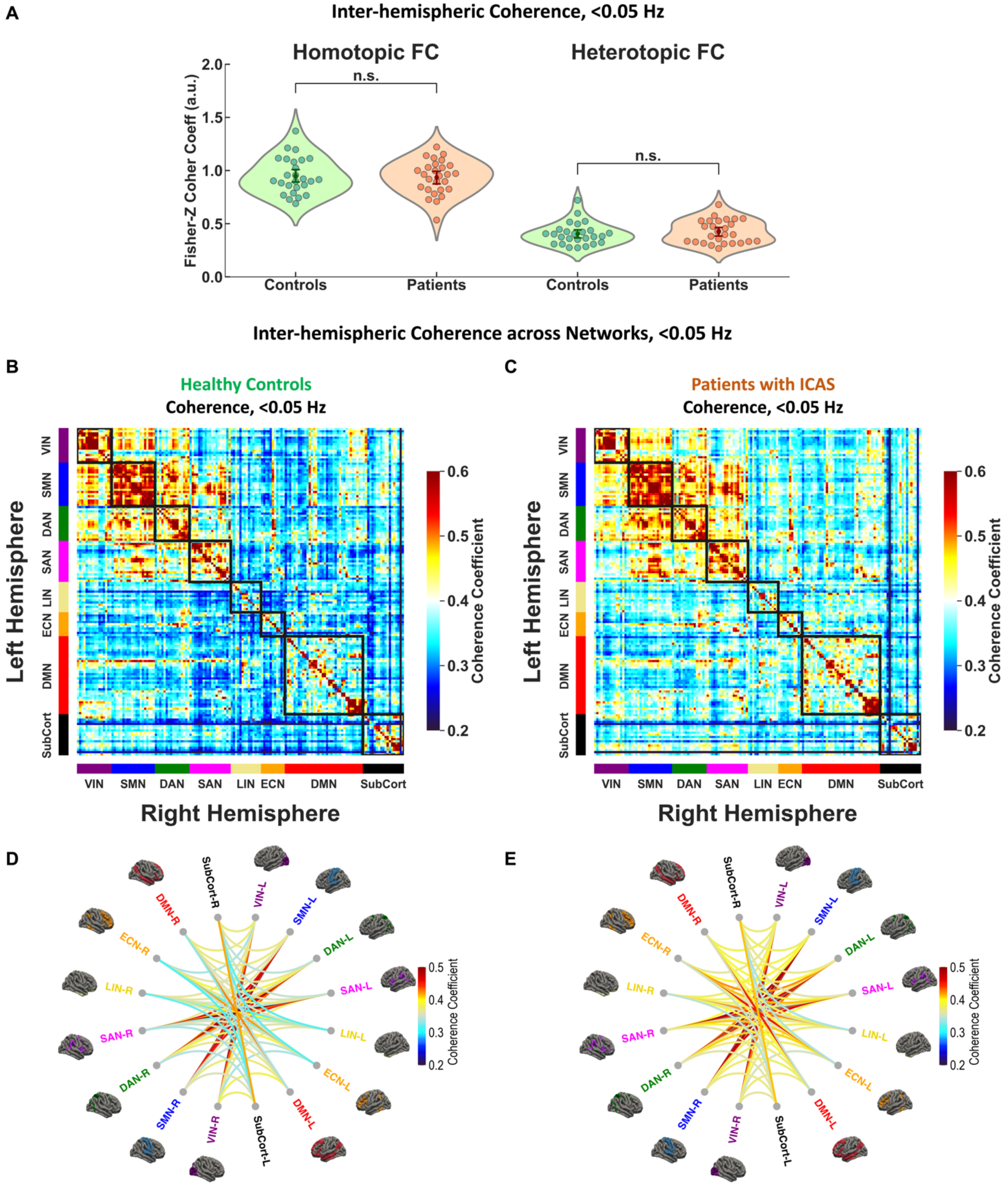

**Figure S3.** Group-level comparisons of inter-hemispheric coherence outside watershed areas at <0.05 Hz. **A.** Inter-hemispheric coherence values for homotopic and heterotopic FCs in healthy controls (green) and patients with ICAS (orange). **B-C.** Network-level inter-hemispheric coherence matrices in healthy controls (**B**) and patients with ICAS (**C**). **D-E.** Corresponding circular visualizations for healthy controls (**D**) and patients with ICAS (**E**). Note that the SAN is not assigned in this analysis.

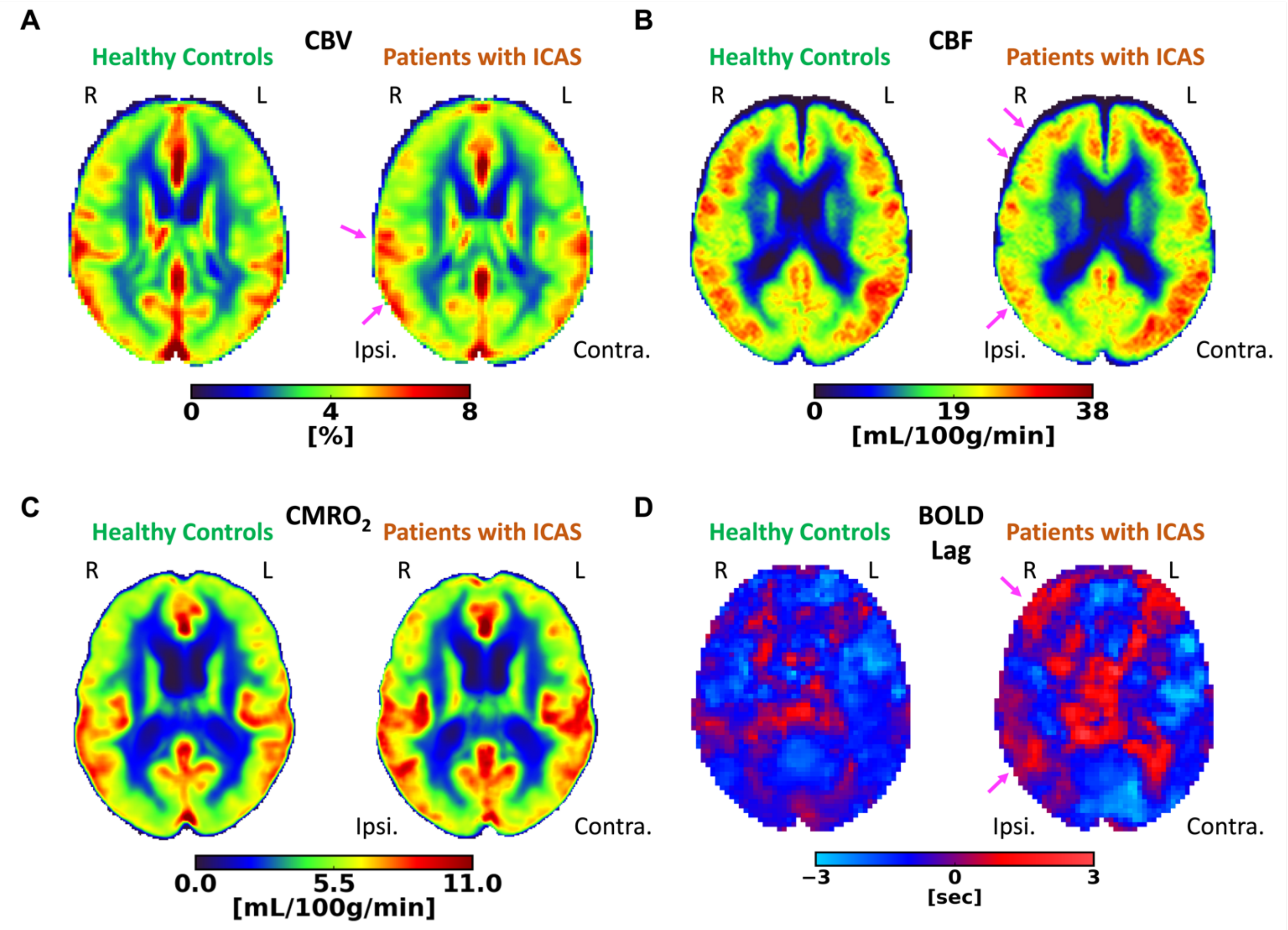


**Figure S4.** Group-averaged multi-parametric quantitative hemodynamic MRI maps in a representative axial slice for healthy controls (left) and patients with ICAS (right). **A**. Relative CBV. **B.** CBF. **C.** Relative $CMRO_2$ **D.** BOLD time lag. 'R' and 'L' denote right and left, respectively. 'Ipsi.' and 'Contra.' indicate hemisphere ipsilateral and contralateral to ICAS. Pink arrows highlight regions with increased lateralization across individual maps.

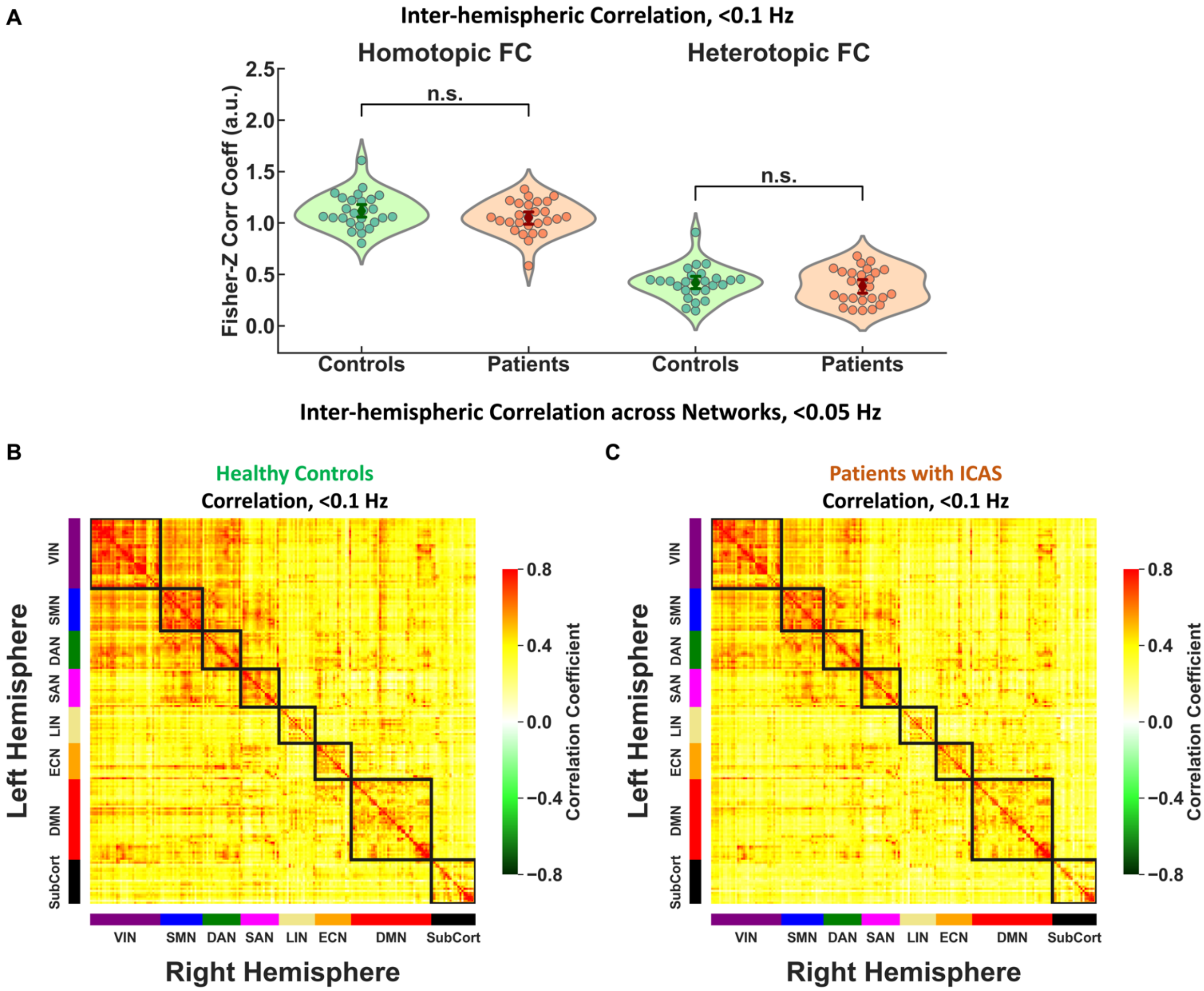


**Figure S5.** Group-level comparisons at <0.1 Hz. **A.** Inter-hemispheric correlation values for homotopic and heterotopic FCs in healthy controls (green) and patients with ICAS (orange). **B-C.** Full network-level correlation matrices in healthy controls (**B**) and patients with ICAS (**C**) across canonical resting-state networks.

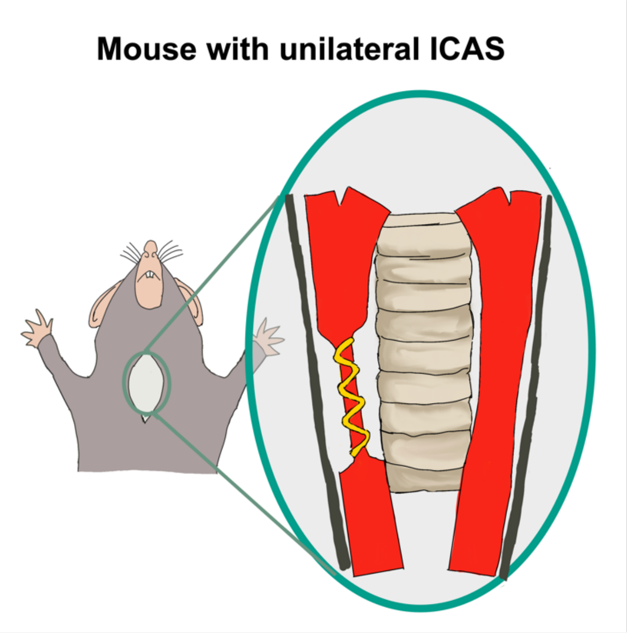


**Figure S6.** Mouse model with  experimentally indiced unliteral ICAS. A submillimeter micro-coil is used to mimic unilateral stenosis.
